# A data-driven approach for timescale decomposition of biochemical reaction networks

**DOI:** 10.1101/2023.08.21.554230

**Authors:** Amir Akbari, Zachary B. Haiman, Bernhard O. Palsson

**Author notes:** Corresponding author. *Email addresses:* (Amir Akbari), (Zachary B. Haiman), (Bernhard O. Palsson).

## Abstract

Understanding the dynamics of biological systems in evolving environments is a challenge due to their scale and complexity. Here, we present a computational framework for timescale decomposition of biochemical reaction networks to distill essential patterns from their intricate dynamics. This approach identifies timescale hierarchies, concentration pools, and coherent structures from time-series data, providing a system-level description of reaction networks at physiologically important timescales. We apply this technique to kinetic models of hypothetical and biological pathways, validating it by reproducing analytically characterized or previously known concentration pools of these pathways. Moreover, by analyzing the timescale hierarchy of the glycolytic pathway, we elucidate the connections between the stoichiometric and dissipative structures of reaction networks and the temporal organization of coherent structures. Specifically, we show that glycolysis is a cofactor driven pathway, the slowest dynamics of which are described by a balance between high-energy phosphate bond and redox trafficking. Overall, this approach provides more biologically interpretable characterizations of network dynamics than large-scale kinetic models, thus facilitating model reduction and personalized medicine applications.

## Introduction

Growth and adaptation are hallmarks of all living systems. They are inherently dynamic processes that are orchestrated by interacting networks of biochemical reactions. Understanding the dynamics underpinning the intricate behavior of biological systems has been a major challenge of systems biology. Kinetic models based on detailed enzymatic rate laws can capture the complex interactions and dynamics of biological system [1, 2]. However, application of these models to large-scale kinetic models is hindered by numerical challenges and their limited interpretability [3].

Biochemical reaction networks have complex structures. Yet, they often exhibit multiscale and comparatively simple dynamics due to the presence of timescale hierarchies [4]. Separation of timescales is believed to be an essential feature of highly evolved reaction networks, conferring stability and robustness [4, 5]. It can lead to the modularization of network dynamics and emergence of ‘independent’ functional units, both of which underlie the robustness and evolvability of living organisms [5, 6]. Although our knowledge of the links between biological complexity and functional modularization is limited, we can gain a deeper understanding of the structural evolution and organization of biochemical reaction networks through systematic studies of their timescale hierarchies.

Timescale decomposition is a common approach for analyzing the dynamics of complex systems. Several techniques have been developed for biochemical reaction networks, most of which fall into two main categories, namely top-down and bottom-up approaches. The first leverages statistical methods and clustering algorithms to identify collections of metabolite with correlated concentration trajectories—referred to as concentration pools—from experimentally measured time-series data [7, 8]. These techniques do not rely on the equations of evolution or kinetic rate laws, although they can use numerically generated time-series data furnished by mass-balance equations. They can handle large-scale systems of varying complexity, but do not mechanistically relate concentration pools to the stoichiometric and dissipative structures of reaction networks. The second determines timescale hierarchies from steady-state eigenvalues [9, 10]. This approach hinges on precise formulations of all reaction rates, providing a mechanistic association between concentration pools and structural properties of reaction networks. Although computations are tractable for largescale networks, inaccuracies may arise for networks with highly nonlinear rate laws where Jacobian spectra are time dependent.

In this paper, we develop a computational technique for timescale decomposition of biochemical reaction networks (Fig. 1), which leverages the strengths of both top-down and bottom-up techniques. This approach, termed Dynamic Mode Analysis (DMA), determines the timescale hierarchy of a reaction network from experimental measured or numerically generated time-series data. It can identify concentration pools for complex networks with uncharacterized rate laws as reliably as top-down techniques. It can also provide a mechanistic description of concentration pools and their organization with respect to the energetics and stoichiometry of reaction networks in the same way as bottom-up techniques. A key component of the proposed technique is an extension of Dynamic Mode Decomposition (DMD) [11]—a technique originally developed to characterize coherent structures arising in fluid flows—that we introduce to identify the dominant exponential decay modes associated with each timescale. We study the timescale hierarchies of hypothetical and biological pathways using DMA, showing that this approach can reproduce the previously characterized concentration pools of these pathways. We also establish a connection between the time-delayed autocorrelation matrix—a statistical descriptor used in top-down techniques [8]—and Jacobian spectra, demonstrating why this descriptor is a useful metric for concentration-pool classifiers.

**Figure 1:**
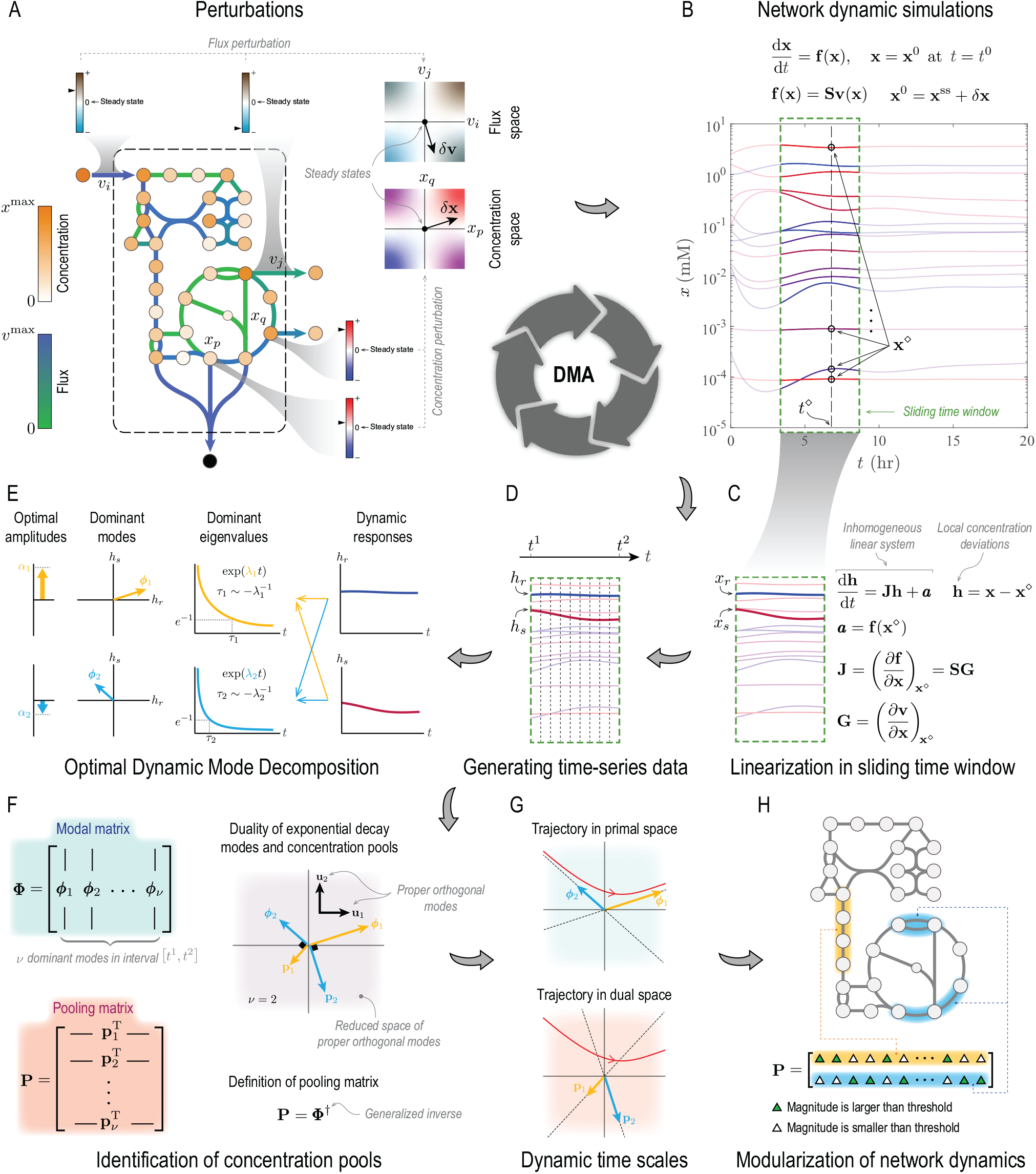
Overview of Dynamic Mode Analysis (DMA). (A) Steady states of biochemical reaction networks are perturbed by introducing concentration or flux disturbances. (B) Concentration trajectories **x**(*t*) are constructed by integrating transient mass-balance equations. (C) For a general nonlinear system, dynamic modes are ascertained by linearizing mass-balance equations in a sliding time window, resulting in a local inhomogeneous linear system. (D) Time-series data are generated by evaluating local concentration deviations **h** at *N* + 1 equally spaced time points in the interval [*t*^1^, *t*^2^] that spans the time window. (E) An Optimal Dynamic Mode Decomposition developed in this work identifies dominant exponential decay modes in the time window from time-series data. (F) The pooling matrix **P**, defined as the Moore-Penrose inverse [12] of the modal matrix **Φ**, is determined from dominant decay modes. (G) Disparate timescales are identified from dominant eigenvalues. (H) Biologically interpretable pools are constructed from the pooling matrix to modularize network dynamics.

## Concentration Pools and Coherent Structures

We first introduce the concept of pools and coherent structures for a case study, providing a biological motivation for their definitions. We define these concept formally in the general case in the next section.

Consider a reaction network involving a single linear pathway with four metabolite and five reactions. We refer to this reaction network as Toy Model 1 (Fig. 2A). The pathway imports Metabolite 1 from and exports Metabolite 4 into the extracellular environment with 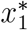 and 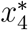 the respective extracellular concentrations. The rate constants of consecutive reactions in this pathway are separated by two orders-of-magnitude (Fig. 2C), so its dynamics are expected to unfold over separate timescales. Dynamic responses to perturbations of steady states are ascertained from transient mass-balance equations

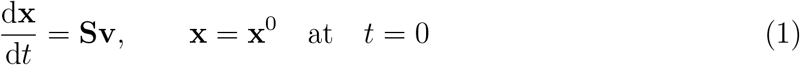

with **x** the concentration vector, **x**^0^ initial concentration perturbations, **S** stoichiometry matrix, x flux vector, and *t* time. Upon perturbations, the dynamics of this system relax over four time intervals associated with three separate timescales *T*_1–3_ (Fig. 2B, dashed lines). Because of the disparity of rate constants, Reactions 1–4 reach their steady states consecutively on their respective timescales. As the dynamics of a given reaction relax, the concentrations of its substrates and products become correlated in the intervals between consecutive timescales. Consequently, trajectories move in a low-dimensional space, suggesting that the dynamics can be adequately described with respect to metabolite pools with correlated concentrations. This is a general behavior that most biochemical reaction networks exhibit, and it is the basis of the definition of concentration pools and coherent structures in this section.

**Figure 2:**
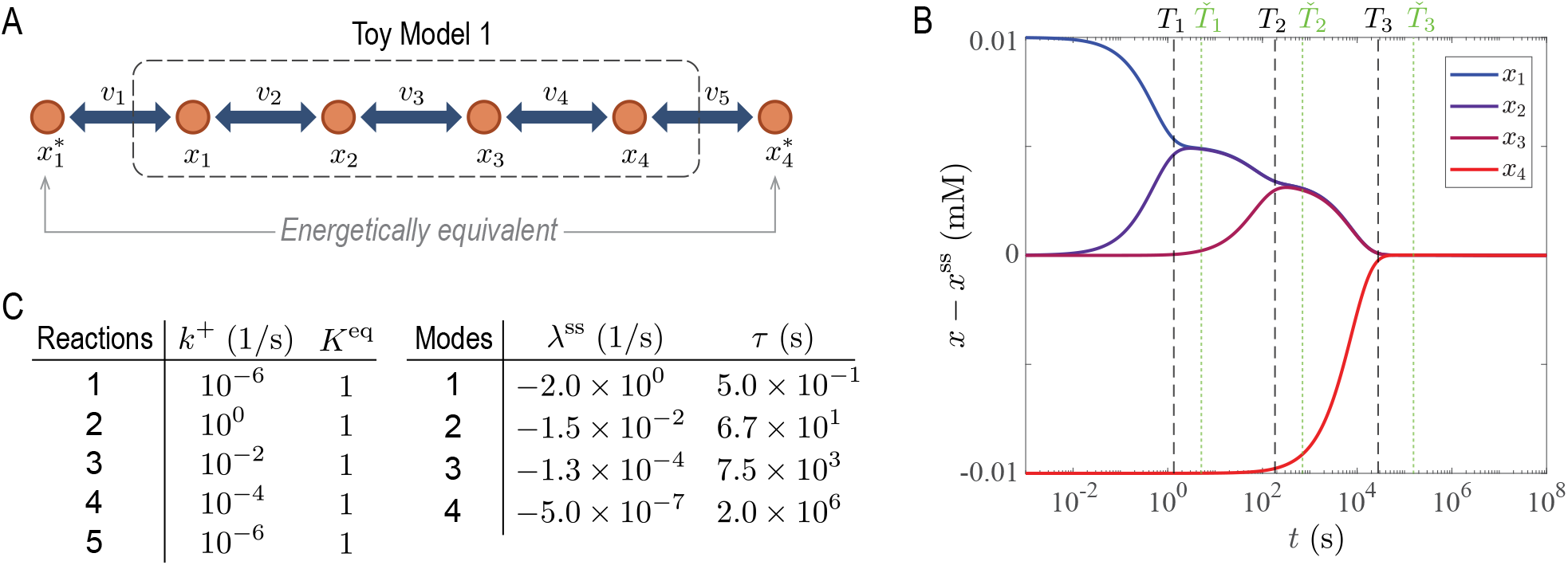
A toy model without energy coupling, where the substrate (Metabolite 1) and product (Metabolite 4) are energetically equivalent. (A) Network map. (B) Dynamic response to concentration perturbations with four characteristic timescales. (C) Rate constants and timescales. The superscript ^*∗*^ indicates metabolite concentration in the extracellular environment.

To formalize the concept of concentration pools, it is more convenient to express concentrations and fluxes relative to their steady-state values. Concentration and flux deviations are defined as ***χ*** := **x −x**^ss^ and ***ϑ*** := **v−v**^ss^, respectively. Accordingly, mass-balance equations can be expressed with respect to these deviation variables

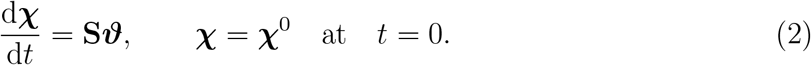

Assuming mass-action kinetics, reaction rates are expressed with respect to deviation variables as

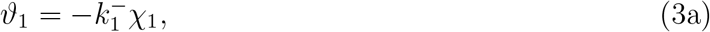

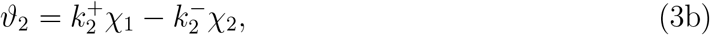

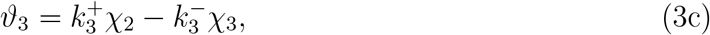

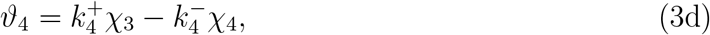

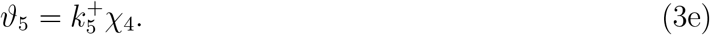

The first pool of Toy Model 1 is associated with the first timescale 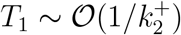 with *𝒪* the order-of-magnitude operator. On this timescale, all flux deviations but *ϑ*_2_ are negligible. Therefore, *ϑ*_2_ *≫ ϑ*_1_, *ϑ*_3_, *ϑ*_4_, *ϑ*_5_, and the mass-balance equations describing the dynamics are

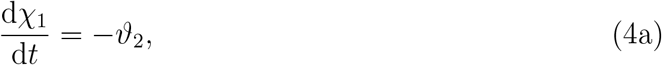

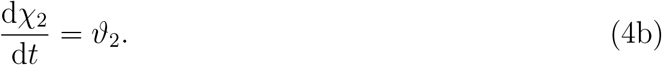

Eliminating *χ*_1_ and *χ*_2_, Eq. (4) can be expressed with respect to *ϑ*_2_ as

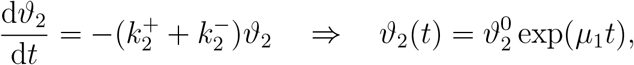

where 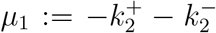, and 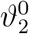 is the initial condition for the flux deviation of Reaction 2. From this analysis, we derive a more accurate approximation of the first timescale as *T*_1_ ∼−1*/µ*_1_. Substituting the solution of *ϑ*_2_ back in Eq. (4), we obtain the trajectory of concentration deviations

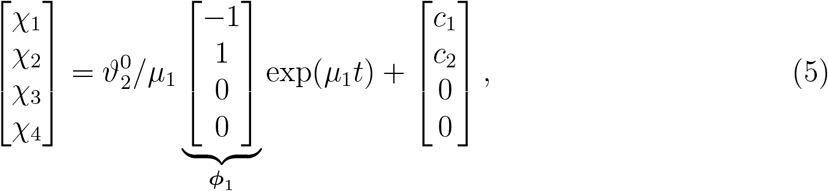

where *ϕ*_1_ is the eigenmode associated with the first pool in the primal space (see Fig. 1F) with *c*_1_ and *c*_2_ the integration constants of Eq. (4). We observe from Eq. (4) that *χ*_1_(*t*) and *χ*_2_(*t*) are negatively correlated on this timescale. Once the dynamics of Reaction 2 have relaxed, its flux deviation equilibrates so that

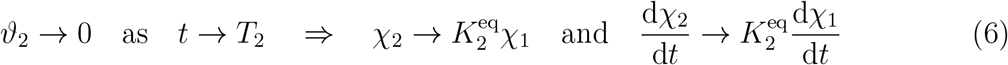

with 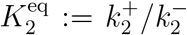 the equilibrium constant of Reaction 2 and *T*_2_ the second timescale. Note that equilibration here refers to the relaxation of flux disturbances as *ϑ*_2_ *→* 0, which is not the same as the equilibration of Reaction 2 when *v*_2_ →0.

If the relaxation time of Reaction 2 is faster than the second timescale of this system, then there is an intermediate timescale *Ť*_1_ ∈ [*T*_1_, *T*_2_] that characterizes the transition between the first and second timescales during which Reaction 2 equilibrates (see Fig. 2B). In this transition period, the flux deviations of Reaction 2 and 3 are of the same order, so that *ϑ*_2_ ∼*ϑ*_3_ ≫*ϑ*_1_, *ϑ*_4_, *ϑ*_5_. To quantify this transitory period, suppose that *ϑ*_2_ = *θϑ*_3_ with *θ* a coefficient that characterizes the transition between the two timescales when *θ ∼ 𝒪* (1). Accordingly, the mass-balance equations simplify to

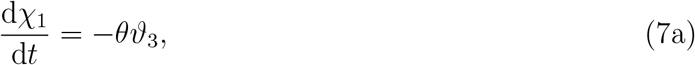

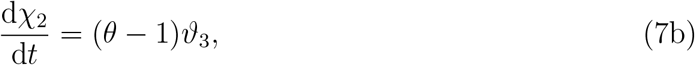

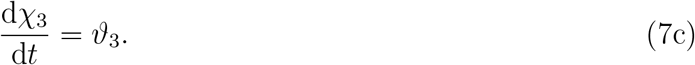

From the asymptotic relation between d*χ*_2_*/*d*t* and d*χ*_1_*/*d*t* stated in Eq. (6), we find that Eq. (7) with 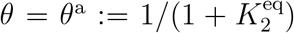 approximates the dynamics in this transitory regime when the second timescale of the system is approached. Note that, in the transitory interval, the coefficient *θ* in these equations varies from a large value *θ ≫*1 near the first timescale to its asymptotic value *θ*^a^ near the second timescale. However, we treat it as a constant to approximate local solutions of Eq. (7) in the transitory regime. We can eliminate *χ*_2_ and *χ*_3_ in Eqs. (7b) and (7c) in the same way as in Eq. (4) to express the mass-balance equations with respect to *ϑ*_3_

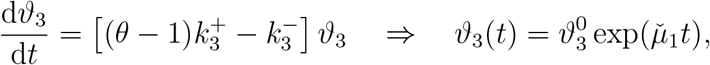

where 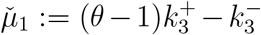, and 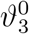 is the initial condition for the flux deviation of Reaction 3. The timescale associated with this regime is estimated 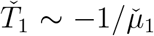, and the trajectory of concentration deviations is ascertained by integrating Eq. (7) using the solution of *ϑ*_3_

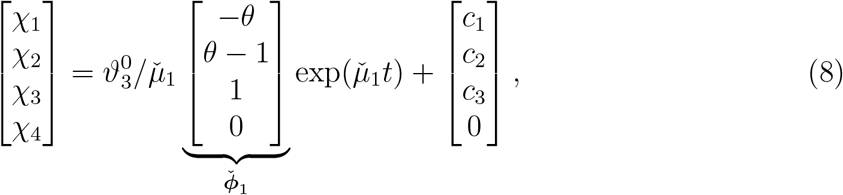

where *c*_1–3_ are integration constants. Note that 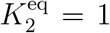 (Fig. 2C) and *θ*^a^ = 1*/*2 in Toy Model 1, implying that *χ*_1_(*t*) and *χ*_2_(*t*) are positively correlated, and together they are negatively correlated with *χ*_3_(*t*) on this timescale. Finally, we highlight an important feature of dominant decay modes during the transition between two consecutive timescales in relation to the role of kinetic parameters. Consider the timescales *T*_1_ and *Ť*_1_ of Toy Model 1 for example. Here, the coefficients of *χ*_1_ and *χ*_2_ in the decay mode *ϕ*_1_ only depend on their stoichiometric coefficients in Reaction 2. However, in the decay mode 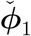, they depend on both the stoichiometric coefficients and rate constants 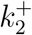 and 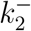.

Having qualitatively described how dynamic trajectories become correlated on the first timescale, we provide a quantitative representation of the first pool. In this work, we define a pool associated with a given timescale as a linear combination of concentrations the time-dependent representation of which aligns with the corresponding exponential decay. If multiple dominant timescales coexist in a time interval, we require the corresponding pools to be dynamically independent. For example, in the transitory period outlined above where both exponential decay modes in Eqs. (5) and (8) are dominant, the first pool *p*_1_ and its transitory counterpart 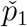 are defined as

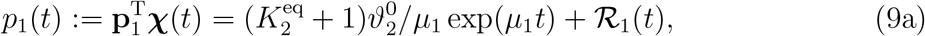

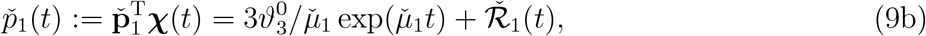

where

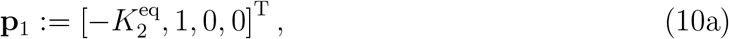

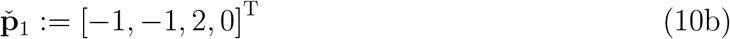

are normalized vector representation of the pools. On occasion, we also refer to this vector representations as pooling maps. Here, ℛ_1_(*t*) and 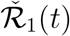 are the residual terms approaching a steady-state value at a faster rate than the exponential decay term for each pool. Moreover, the timescale associated with each pool is defined as a time point beyond which the pool is sufficiently close to its steady state value (Fig. 3A). For example, the timescales of the foregoing two pools are defined as

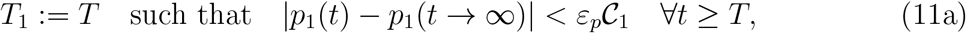

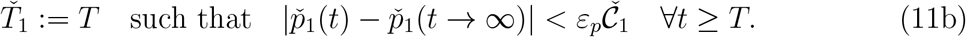

Here, *ε*_*p*_ is a tolerance threshold controlling the closeness of pools to their steady-state values with *𝒞*_1_ and 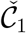 appropriate concentration scales. For example, for *p*_1_ and 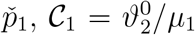 and 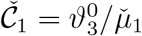 are reasonable concentration scales.

As noted in previous works, a proper definition of pools should ensure that each pool is dynamically independent of other pools in a system when they form on the same timescale [13, 14]. In top down approaches, such as Principal Component Analysis or Independent Component Analysis, orthogonality and independence are equivalent concepts [15]. However, as discussed at the beginning of this section, a biological motivation for introducing the concept of concentration pools is identifying aggregate variables that naturally arise in systems with low-dimensional dynamics—a key characteristic of reaction networks with timescale hierarchies. Accordingly, alignment with exponential decay modes associated with separate timescales is considered a more appropriate notion of independence [11, 13]. Since eigenmodes are not necessarily orthogonal for a general reaction network, dynamical independence does not imply the orthogonality of pools in the concentration space. In this work, we adopt the same notion of independence. However, in addition to alignment with exponential decay modes, we also require a reciprocal orthogonality between the vector representation of the pools and eigenmodes. Here, we present this reciprocal relationship for *p*_1_ and 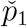 in the transitory regime between the first two timescales of Toy Model 1, deferring the formulation for the general case to the next section. The reciprocal orthogonality conditions for the first pool and its transitory counterpart are

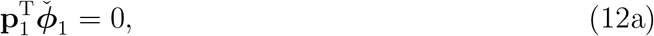

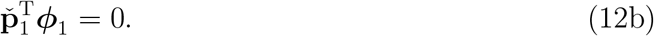

These conditions are motivated by their important geometric and physical interpretations, and they are linked to the concept of flux-concentration duality [16] in chemical reaction networks (see Appendix A and Fig. 1F).

Coherent structure is another concept that we examine in this paper. Its definition, which is derived from but is more restrictive than that of concentration pools, centers around correlations among concentration trajectories. As we noted for Toy Model 1, the dominant eigenmodes on a given timescale span a reduced concentration space in which the dynamics unfold. Accordingly, metabolite coefficients in the pooling maps, which are in turn ascertained from the respective eigenmodes, determine metabolites the concentrations of which are affected on that timescale. The concentration trajectories of these metabolites remain correlated until the next timescale of the system has been reached. For a linear system, such as Toy Model 1, the eigenmodes and their respective pooling maps do not vary with time. Therefore, the correlation coefficients among metabolites that are present in a pool remain constant. However, in a general nonlinear system, the Jacobian matrix and its eigenmodes can vary along dynamic trajectories. Consequently, correlation coefficients also become time-dependent, but they can plateau in small intervals between the timescales of the system.

**Figure 3:**
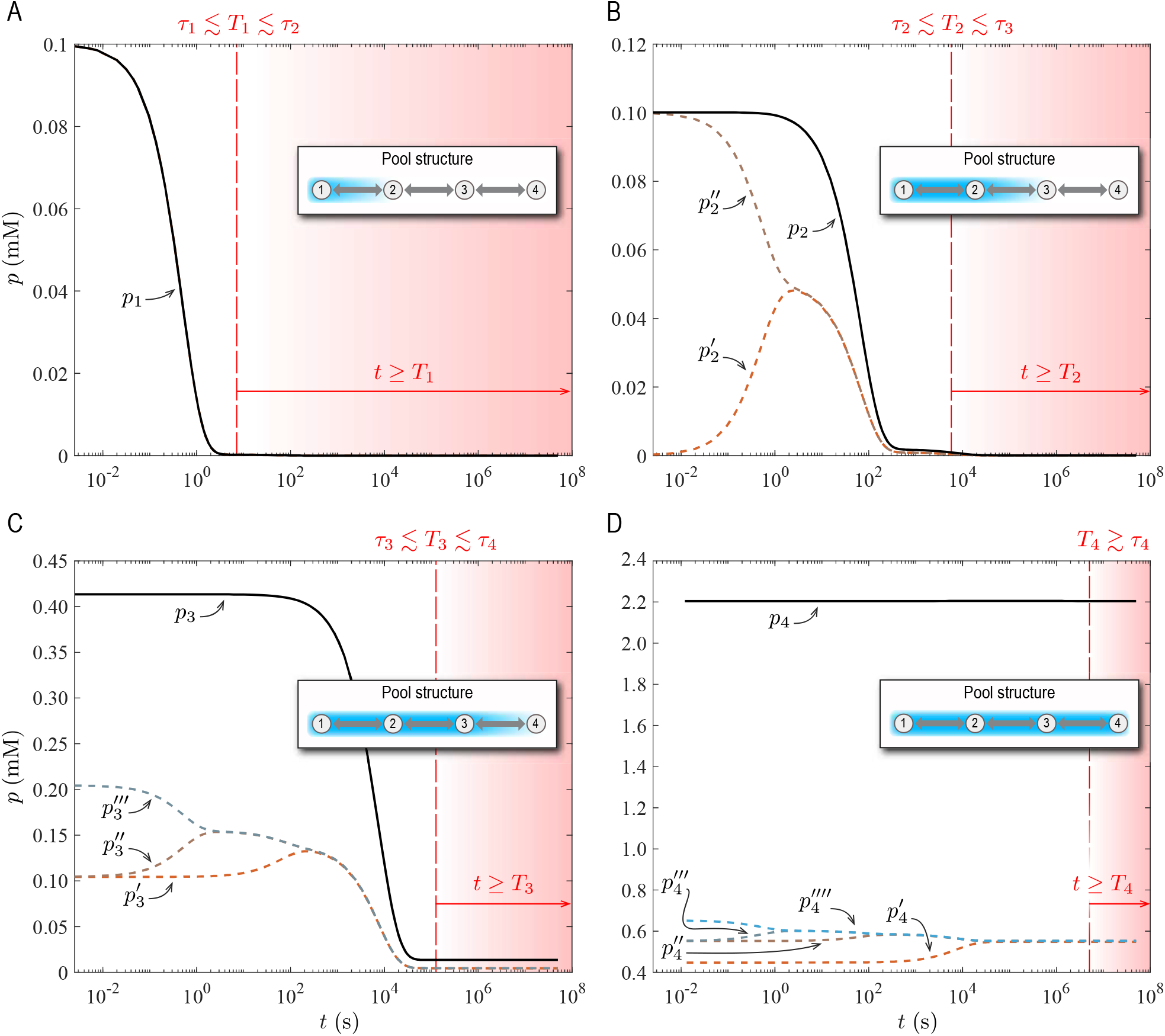
Pool representations for Toy Model 1 described in Fig. 2. (A) Representation of Pool 1 associated with the disequilibrium of Reaction 2. (B) Representations of Pool 2 associated with the disequilibrium of Reaction 3. (C) Representations of Pool 3 associated with the disequilibrium of Reaction 4. (D) Representations of Pool 4 associated with the conservation of all the intracellular metabolites. Timescales *τ*_1–4_ are provided in Fig. 2B. The fourth timescale is defined as *T*_4_ := *Ť*_3_ (see Fig. 2B), where *Ť*_3_ is a timescale characterizing a transitory regime between the third timescale and steady state of Toy Model 1.

For a general nonlinear reaction system, we define a coherent structure as a subset of metabolites in concentration pool the correlation coefficients of which vary within a prescribed tolerance threshold in time intervals between the timescales of the system. To make this definition more precise, suppose that metabolite *i* and *j* in a reaction system are part of a pool that becomes activate in the interval [*T*_1_, *T*_2_], where *T*_1_ and *T*_2_ are two consecutive timescales. Then, for these metabolites to form a coherent structure, their concentration trajectories must satisfy

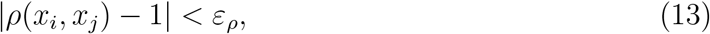

where *ε*_*ρ*_ is a tolerance threshold controlling variations of correlation coefficients, and the correlation function *ρ* is given by

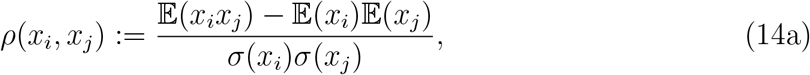

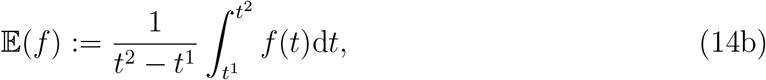

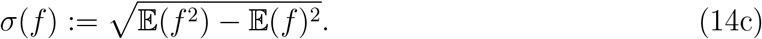

Here, 𝔼(*f*) and *σ*(*f*) are the expectation and standard deviation of a time-dependent function *f* (*t*). Moreover, the correlation function is evaluated in the interval [*t*^1^, *t*^2^] in which the coherent structure forms, where *T*_1_ ≤*t*^1^ *< t*^2^ ≤*T*_2_. Note that superscripts of *t* in Eq. (14) are indices referring to time-interval bounds and should not be confused with exponents. For coherent structures with more than two metabolites, Eq. (13) must be satisfied for all pairwise correlations among the metabolites. For linear systems, such as Toy Model 1, concentration pools always form a coherent structure because the pooling maps and correlation coefficients are constant along dynamic trajectories. Note also that coherent structures can generally span several timescales.

Having discussed the concepts of concentration pools and coherent structures for the first timescale of Toy Model 1, we follow the same procedure to characterize concentration pools and coherent structures that form at slower timescales. The second pool arises when the dynamics of Reaction 3 begin to relax. It is associated with the timescale *T*_2_ *∼ −*1*/µ*_2_ with 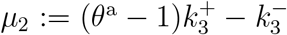. On this timescale, we have *ϑ*_2_ *≃ θ*^a^*ϑ*_3_ *≫ ϑ*_1_, *ϑ*_4_, *ϑ*_5_, leading to the following expression for the second pool

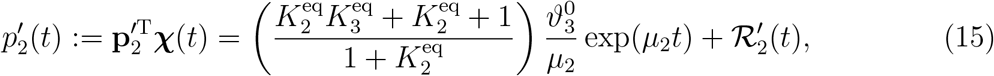

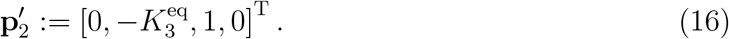

Here,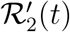 denotes a residual term that approaches a steady-state value at a faster rate than the exponential decay term, so its leading-order term is exp(*µ*_1_*t*). Another expression for the second pool can be derived by combining 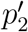 with pools that form on faster timescales than *−*1*/µ*_2_. For example, once the dynamics of Reaction 2 have relaxed in the transitory interval [*Ť*_1_, *T*_2_], *χ*_1_(*t*) and *χ*_2_(*t*) become correlated through the relationship

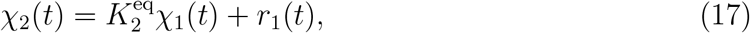

where *r*_1_(*t*) is a residual, the leading-order term of which is exp(*µ*_1_*t*). Substituting Eq. (17) in Eq. (15) results in

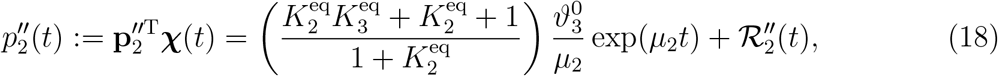

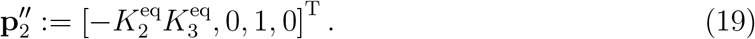

A sum-total pool can also be constructed by adding the first two expressions in Eqs. (15) and (18), resulting in

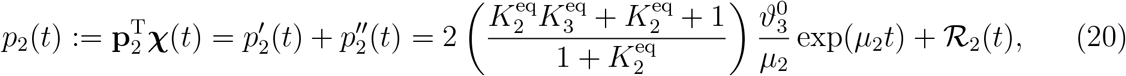

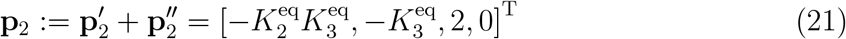

with *ℛ*_2_(*t*) the sum-total residual term. Note that the leading-order terms of 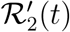,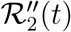, and *ℛ*_2_(*t*) are exp(*µ*_1_*t*). Thus,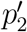,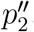, and *p*_2_ can be regarded as three representations of the second pool since they all align with exp(*µ*_2_*t*), exhibiting a similar asymptotic behavior for *t* ≳ *T*_2_ (Fig. 3B). Note that the pooling maps for all three representations are orthogonal to the second transitory mode that becomes active in the interval [*Ť*_2_, *T*_3_], so they satisfy the reciprocal orthogonality condition

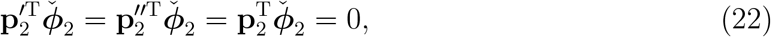

which is similar to the reciprocal orthogonality condition in Eq. (12a) for the first timescale.

The third timescale *T*_3_ is associated with the relaxation of Reaction 4. The procedure for characterizing its concentration pool is similar to that outlined for the second timescale and will not be repeated here (see Appendix A2 for details) . We only highlight the representations of the third pooling map

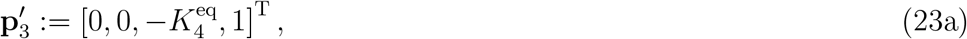

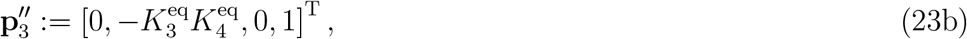

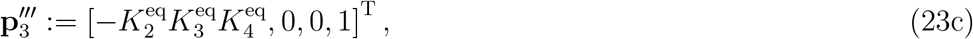

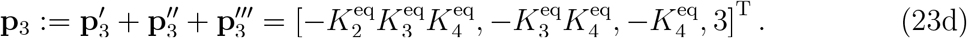

The corresponding pools 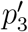,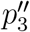,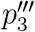, and *p*_3_ exhibit a similar asymptotic behavior for *t* ≳ *T*_3_ (Fig. 3C). As with the second timescale, **p**_3_ denotes the sum-total representation of the third pooling map.

The dynamics of Reaction 4 relax fully once the initial perturbations have propagated throughout the network and reached the boundary reactions. Upon relaxation, all flux and concentration disturbances equilibrate towards a steady state on a transitory timescale *Ť*_3_. One this timescale, which we regard as the fourth timescale of Toy Model 1 (*i*.*e*., *T*_4_ := *Ť*_3_), all the concentration deviations align with the slowest eigenmode, forming a concentration pool containing all the intracellular metabolites. Since this pool corresponds to the only dominant eigenmode for *t* ≳ *T*_4_, there are no slower eigenmodes to evolve into; hence, it needs not satisfy any reciprocal orthogonality conditions. Accordingly, the concentration pool associated with the largest timescale is defined as a normalized form of the slowest eigenmode. This condition automatically arises from the general definition of the pooling matrix—a matrix containing all the dominant pooling maps on a given timescale—which will be formalized for a general nonlinear reaction system in the next section.

The fourth pool has the following representations (see Appendix A2 for details)

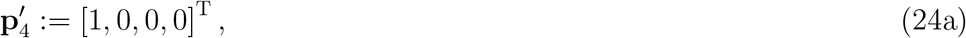

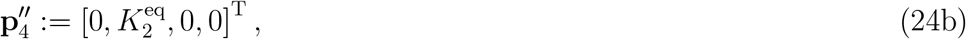

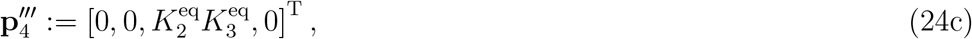

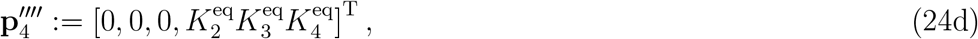

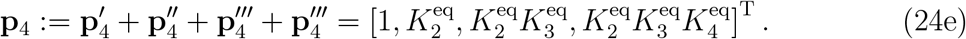

The asymptotic profiles of the respective pools 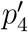,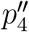,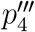,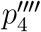,and *p*_4_ are similar for *t* ≳ *T*_4_ (Fig. 3D). As before, **p**_4_ denotes a sum-total representation.

Lastly, we emphasize that the purpose of the analysis presented here and Appendix A2 is to elucidate the connections between concentration pools, timescales, and the equilibration of flux disturbances using approximate methods. This approach is justified since a fundamental characteristic of biochemical reaction networks underlying the separation of timescales is the presence of vastly disparate rate constants. The solutions provided in this section and Appendix A2 are not exact, so the eigenvalues *µ*_*i*_ estimated here should be regarded as approximations of the exact eigenvalues *λ*_*i*_. Since Toy Model 1 is a linear system, its eigenvalues remain constant along dynamic trajectories. Consequently, in any time interval, the exact solution of mass-balance equations can be expressed as a linear combination of exponential decay modes with varying amplitudes without needing transitory eigenvalues. However, the reason why we encountered the transitory eigenvalues 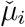 here lies in our approximation method describing the kinetics on a given timescale by the flux of a single reaction the rate constant of which gives rise to that timescale.

## Dynamic Mode Analysis

DMA is a data-driven approach to identify timescale hierarchies of a chemical reaction network and characterize its dynamic responses to flux or concentration perturbations (Fig. 1). Specifically, this approach aims to compute concentration pools and coherent structures that emerge as a reaction network evolve in time from time-series data. Although data can be generated from experimental measurements or numerical simulations, we only focus on numerical solutions of kinetic models of biochemical reaction networks in this paper. We assume that the concentration trajectories of all the metabolite in the network of interest are given. The goal is then to identify the dominant eigenvalues, dominant eigenmodes, and the respective concentration pools algorithmically in any time interval along dynamic trajectories of the network.

The algorithm begins by solving the mass-balance equations Eq. (1) for a reaction network with *n* metabolites and *m* reactions, the steady-state concentrations or fluxes of which is perturbed (Fig. 1A). The resulting concentration trajectory **x**(*t*) is then computed numerically (Fig. 1B). The goal in subsequent steps of DMA is to identify the dominant eigenvalues and eigenmodes in a sliding time window spanning the interval [*t*^1^, *t*^2^] as it moves from *t* = 0 to *t* = *T*_*∞*_, where *t* = *T*_*∞*_ is the total relaxation time—a time by which all flux and concentration disturbances have relaxed to within a tolerable threshold. To analyze the dynamics in the sliding time window, the mass-balance equations are linearized locally around a reference time *t*^*⋄*^ *∈* [*t*^1^, *t*^2^]

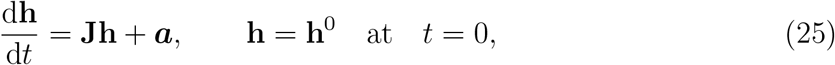

where

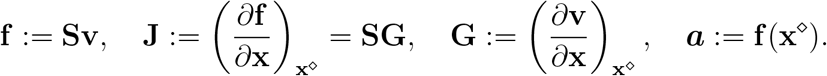

Here, **J** is the local Jacobian matrix, **x**^*⋄*^ := **x**(*t*^*⋄*^) the concentration vector at the reference time, and **h** := **x −x**^*⋄*^ the local concentration deviations. Note that, for a general nonlinear reaction network, Eq. (25) is an inhomogeneous system of equations, the solutions of which cannot be expressed with respect to purely exponential decay terms (Fig. 1C). Thus, we recast it into the homogenous system

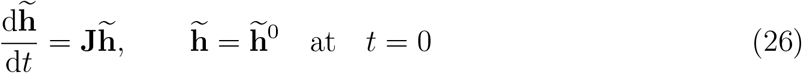

by introducing a new variable 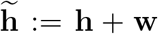, we term the local transformed concentration deviation, where **w** := **J**^*−*1^a captures the inhomogeneity of Eq. (25). The general solution of Eq. (26) is written [17]

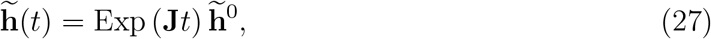

where Exp is the exponential map. Since we are only concentrated with small perturbations of noncritical stable steady states, we assume that *ℜ𝔢*(λ) *<* **0** along all dynamic trajectories. Therefore, **J** is always non-singular, so **w** is a well-defined quantity. Here, *ℜ𝔢*(*·*) returns the real part of a complex argument, and λ the eigenvalue vector of the Jacobian. The linearization of mass-balance equations introduced here allows the application of top-down timescale decomposition techniques (*e*.*g*., DMD) to identify timescale hierarchies from time-series data.

In the next step, time-series data are generated from the numerical solution of local concentration deviations in the current time window by evaluating **h**(*t*) at *N* + 1 equally spaced time points in [*t*^1^, *t*^2^] (Fig. 1D), compiling their values in two data matrices

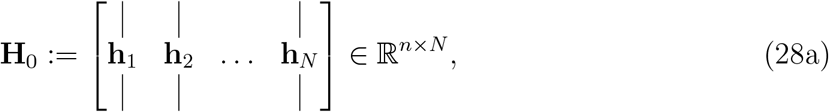

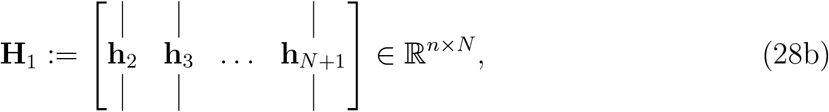

where **h**_*k*_ := **h**(*t*_*k*_) is the vector of local concentration deviations evaluated at the *k*th time point in [*t*^1^, *t*^2^] with Δ*t* := (*t*^2^ *− t*^1^)*/*(*N −* 1) the gap between consecutive time points. The corresponding transformed data matrices 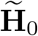 and 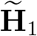 are similarly defined in terms of 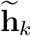. From Eq. (27), we find the relationship between transformed concentration deviations at two consecutive time points

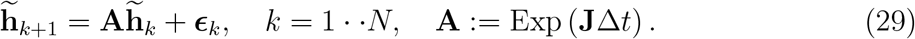

Here, ***ϵ***_*k*_ is an error vector associated with the linearization of mass-balance equations. It also accounts for the errors arising from numerical simulation (*e*.*g*., truncation errors) or experimental measures depending on how data is generated. Since **A** is a constant matrix, we can write Eq. (29) in a matrix form

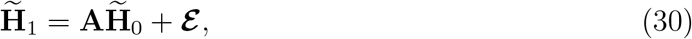

where ***ℰ*** is a matrix of the same size as 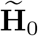 and 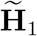, containing all the error vectors ***ϵ***_*k*_.

If mass-balance equations are known, or there are approximation methods to estimate **w** from time-series data, then the dominant eigenvalues and eigenmodes of the reaction network in the current time window can be directly determined from 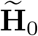 and 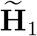 using top-down approaches, such as DMD [11] or Optimal Dynamic Mode Decomposition (ODMD) [18, 19]. In the following, we briefly outline the procedure for ODMD.

To determine the dynamic dimensionality of the system, the singular-value decomposition of 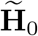 is computed

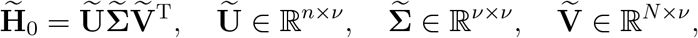

where *v* is the number of singular values that are nonzero to within a tolerable threshold *ε*_SVD_. The columns of Ũ are the proper orthogonal modes [19] of 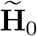, and they span a reduced *v*-dimensional subspace of the local transformed concentration-deviation space where the dynamic trajectories lie in the current time window. We refer to this subspace as the space of transformed proper orthogonal modes. The projection of **A** onto this reduced space is

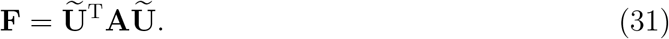

The projected matrix **F** ∈ ℝ^*ν×ν*^ in this equation can be proved to be an upper Hessenberg matrix [20]. Because **F** and **A** are related through a similarity transformation, **F** contains the dominant eigenvalues of **A** in the current time window. Errors in Eq. (30) are then minimized by solving the optimization problem

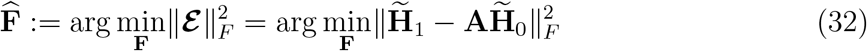

to identify a matrix 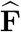 that best describes the data in 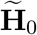 and 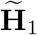, where ∥·∥_*F*_ denotes the Frobenius norm. Substituting **A** from Eq. (31) in Eq. (32), the solution of the foregoing optimization problem is ascertained [19]

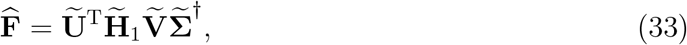

where the superscript ^*†*^ denotes the Moore-Penrose generalized inverse [12]. The eigenmodes of 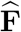 are the projections of the dominant eigenmodes of **A** in the space of transformed proper orthogonal modes, and it eigenvalues are the dominant eigenvalues of **A**.

In this paper, both **F** and **w** are treated as unknowns, so 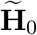 and 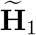are not provided at the outset. Therefore, in the following, we introduce an extension of ODMD to determine the dominant eigenvalues and eigenmodes of the inhomogeneous system Eq. (25) directly from the time-series data in **H**_0_ and **H**_1_. We start by rewriting Eqs. (29) and (30) in terms of local concentration deviations

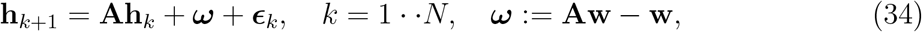

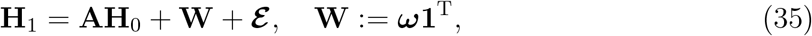

where **1** is an all-one column vector with *N* components. As previously stated, because of the inhomogeneous term ***ω*** in Eq. (34) or **W** in Eq. (35), DMD or ODMD are not directly applicable, although we can show that ***ω*** →**0** in the limit Δ*t →*0 (see Appendix A3). Thus, the inhomogeneity can theoretically be eliminated from Eq. (34) by generating arbitrarily fine temporal grid in the sliding time window. However, it is impractical to do so because of the computational costs and additional errors it introduces in subsequent steps of the algorithm. Following the same procedure as DMD, the singular-value decomposition of **H**_0_ is computed

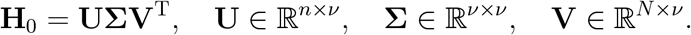

As in the previous case, *v* reflects the dynamic dimensionality of the system in the current time window, and the columns of **U** are the proper orthogonal modes of **H**_0_, spanning the local concentration-deviation space. We refer to this subspace as the space of proper orthogonal modes. As before, **A** is expressed with respect to the proper orthogonal modes through the similarity transformation

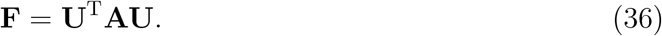

The goal now is to find **F** and ***ω*** such that the errors in Eq. (35) are minimized. The corresponding optimization problem is written

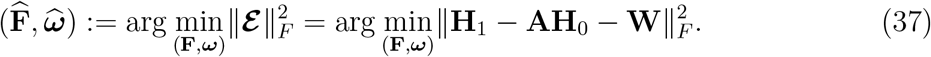

This is an unconstrained quadratic program, so its solution can be determined analytically (see Appendix A4 for the proof)

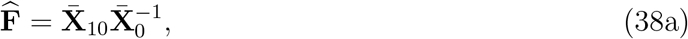

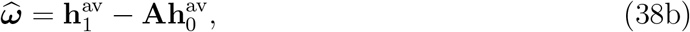

where

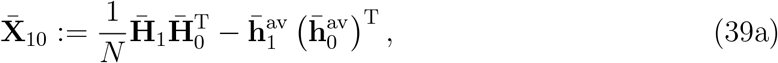

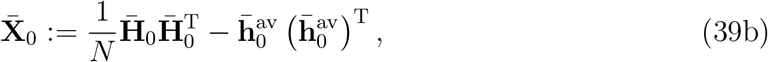

and

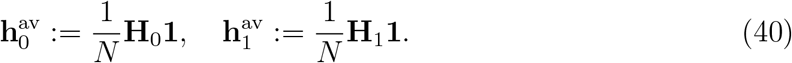

Note that the barred vectors and matrices denote the representation of their unbarred counterparts in the space of proper orthogonal modes. Accordingly,

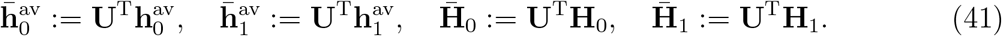

Here,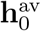 and 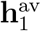the average of concentration-deviation data in **H**_0_ and **H**_1_, respectively. Moreover,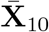 and 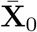 are the time-delayed autocorrelation and covariance matrices of the time-series data represented with respect to the proper orthogonal modes. Note that Eq. (38a) may be regarded as a generalization of the DMD solution Eq. (33) it approaches to as ***ω*** →**0**, which in turn occurs when the steady state has been attained. We also emphasize that metrics based on time-delayed autocorrelation were used in previous top-down approaches along with clustering algorithms to identify concentration pools irrespective of the underlying biochemical mechanisms [7, 8]. However, as we highlighted in the previous section, what underlies the of formation concentration pools and coherent structures in biochemical reaction networks is the presence of timescale hierarchies. Furthermore, the optimal solution 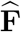 in Eq. (38a) implies that the time-delayed autocorrelation matrix contains information about the local Jacobian spectra. Although the expressions used in previous studies are not identical to Eq. (38a), our analysis demonstrate why time-delayed autocorrelation is a useful metric for statistical classifications of concentration pools.

Once 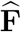 has been computed from Eq. (38a), the dominant eigenvalues and eigenmodes can readily be determined (Fig. 1E). Let 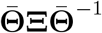 be the eigenvalue decomposition of 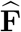 with **Ξ** a diagonal matrix, where the diagonal entries Ξ_*ii*_ := *µ*_*i*_ are the eigenvalues of 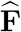 (not to be confused with the approximate eigenvalues of Toy Model 1 in the previous section). Then, the columns of 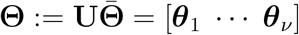 correspond to the dominant eigenmodes of **A** in the current time window. Because **A** and **F** are related through a similarity transformation, *µ*_*i*_ are also the dominant eigenvalues of **A**. From Eq. (29), it follows that

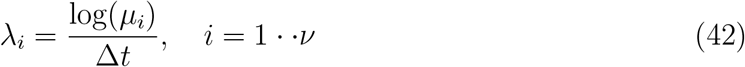

with *λ*_*i*_ the dominant eigenvalues of the local Jacobian matrix **J**.

The next steps of DMA are similar to those of ODMD [19], but modifications are required due to the inhomogeneity of Eq. (34). Here, we note that the inhomogeneous term ***ω*** can be eliminated by introducing the differential time-series data matrix **D** := **H**_1_ *−* **H**_0_, so that

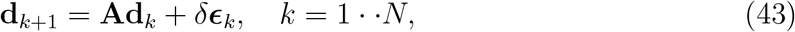

where **d**_*k*_ := **h**_*k*+1_ −**h**_*k*_ is the *k*th column of **D** and *δ****ϵ***_*k*_ := ***ϵ***_*k*+1_ − ***ϵ***_*k*_. Since Eq. (43) is a homogenous system, we can follow the same procedure as ODMD to determine the optimal amplitudes associated with the dominant eigenvalues and eigenmodes ascertained in the previous step. Let

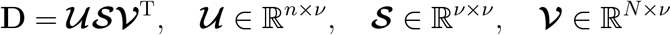

be the singular-value decomposition of **D** ∈ ℝ^*n×N*^ and **Γ** ∈ ℂ^*ν×N*^ the Vandermonde matrix constructed from the eigenvalues {*µ*_1_, *· · ·, µ*_*ν*_}. Then, the optimal amplitudes α are given by [19]

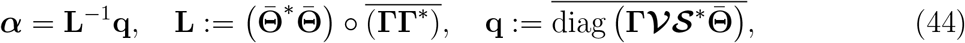

where an overline denotes the complex-conjugate of a matrix, the superscript ^*∗*^ denotes the complex-conjugate transpose, and ∘ indicates elementwise matrix multiplication. Accordingly, the modal matrix reads

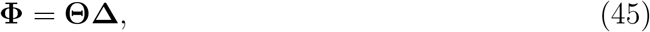

the columns of which *ϕ*_*i*_ are the dominant eigenmodes of **A**. Here, **Δ** ∈ ℂ^*ν×ν*^ is a diagonal matrix with diagonal entries Δ_*ii*_ := *α*_*i*_. Finally, we define the pooling matrix as

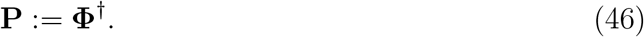

Note that the pooling maps defined in the previous section correspond to the rows of **P**. Moreover, the definition Eq. (46) ensures that the modal and pooling matrices satisfy the reciprocal orthogonality conditions in all time intervals. It is also consistent with the relaxation of the slowest eigenmode discussed in the previous section. Recall, we defined the pooling map near the steady state corresponding to the slowest mode as a normalized form of the slowest eigenmode, which is compatible with Eq. (46) since the Moore-Penrose inverse of a vector yields the transpose of same vector with a normalized length.

## Results

We illustrate the application of DMA by analyzing the timescale hierarchies of chemical reaction networks in three case studies. The first two are hypothetical pathways that possess key characteristics of typical biological pathways. The last is glycolysis—a well-studied biological pathway the concentration pools of which were previously characterized [9, 13]. We examine this case study to validate our computational framework by reproducing some of the known concentration pools of glycolysis and their respective timescales.

### Toy Model 1

The first case study is Toy Model 1, which is the same pathway we examined in previous sections (see Figs. 2 and 3). It converts a substrate (Metabolite 1) into a product (Metabolite 2) without energy coupling. The substrate and product are energetically equivalent, and all the equilibrium constants are of the same order-of-magnitude 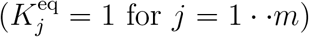 therefore, the flux is driven through the pathway by maintaining a gradient between the extracellular concentrations 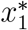 and 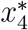 (Fig. 4A). The boundary reactions (Reactions 1 and 5) are the rate-limiting steps, and the rate constants of consecutive intracellular reactions are separated by two orders-of-magnitude (Fig. 2C).

**Figure 4:**
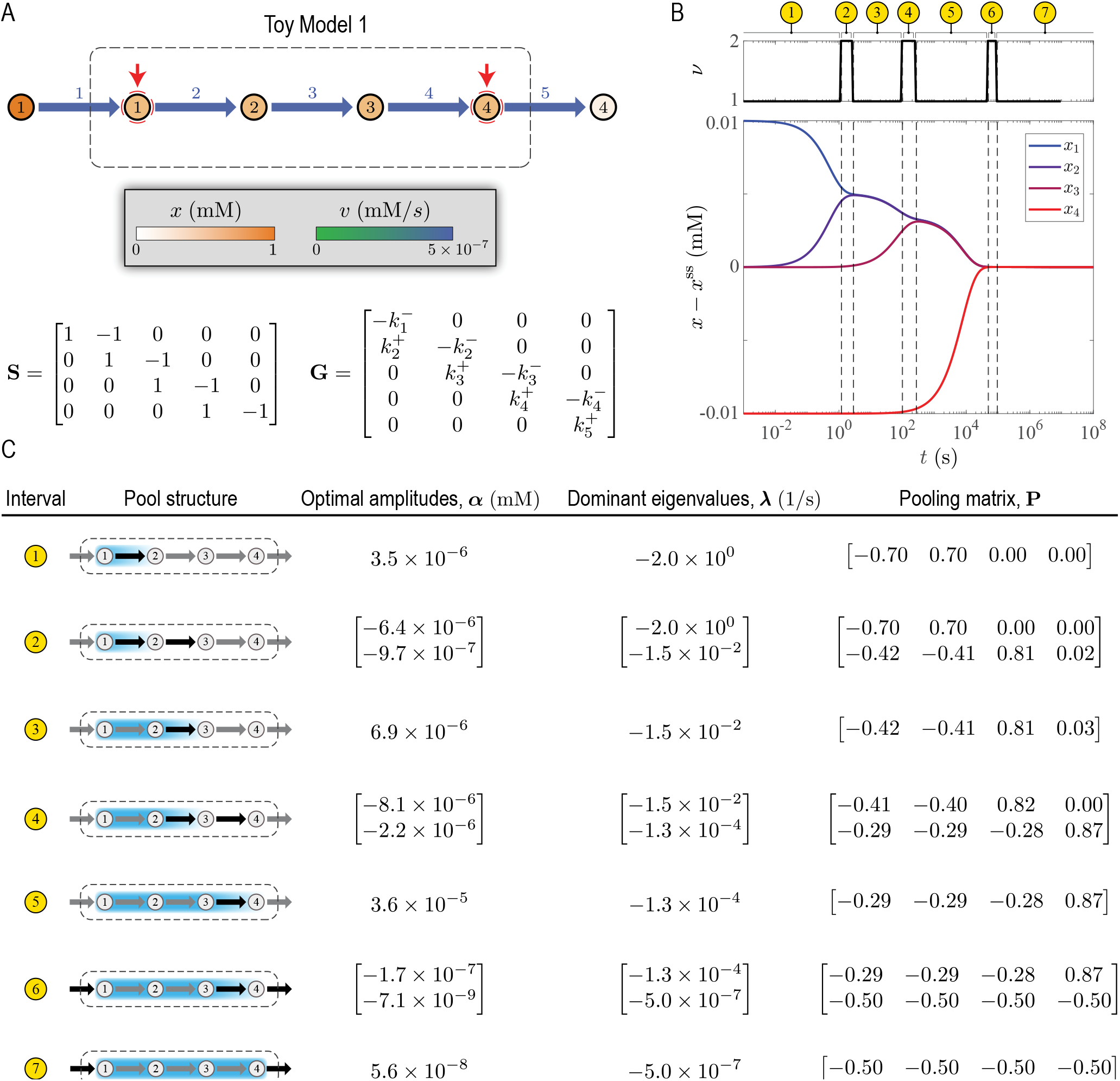
Dynamic Mode Analysis of Toy Model 1 described in Fig. 2. (A) Visualization of the steady state solution in a network map. Red arrows indicate metabolites the concentration of which is perturbed. (B) Dynamic response to concentration perturbations. Deviations from the steady state *x− x*^ss^ and dimensionality *ν* are plotted along the dynamic trajectory. Dashed lines indicate a transition between two time periods with distinct exponential decay modes, partitioning the overall relaxation time into seven characteristic time intervals. (C) Optimal amplitudes, dominant eigenvalues, and pooling matrix for each time interval identified in (B). Black arrows in pool structures indicate parts of the network that are affected the most by the initial perturbation in the corresponding time interval. Gradient and uniformly colored envelops enclose metabolites the concentration trajectory of which are negatively and positively correlated in the corresponding time interval, respectively.

Toy Model 1 has a timescale hierarchy due to the separation of rate constants. We identified its timescales and their respective concentration pools using DMA (Fig. 4). We induced a dynamic response by perturbing *x*_1_ and *x*_4_, which activated all three timescales associated with the rate constants of the intracellular reactions (Fig. 4A,B). The successive equilibration of the intracellular flux disturbances partitioned the total relaxation time into seven distinct time intervals, corresponding to the three main timescales and their transitory counterparts we highlighted previously (Fig. 4B,C). Of particular interest are the slowest transients near the steady state. Here, all the intracellular metabolites pool together into a single aggregate metabolite with coefficients (1, 1, 1, 1), the concentration of which is controlled by the boundary reactions. Overall, the timescales and pooling matrices furnished by DMA for each time interval agreed well with our estimates based on the approximate analytical method presented in Section “Concentration Pools and Coherent Structures” (Fig. 4C).

Furthermore, DMA could accurately characterize the transitory regime between two consecutive timescales, which we discussed earlier. DMA identified two stages associated with the relaxation of each reaction. In the first, the concentrations evolve in a direction corresponding to a maximal energy dissipation of the reaction, where the substrate and product concentrations are negatively correlated (*e*.*g*., see the coefficients of *x*_1_ and *x*_2_ in the first pooling map of Interval 2). In the second, the concentrations evolve in a direction associated with the equilibration of the flux disturbance, where the substrate and product concentrations are positively correlated (*e*.*g*., see the coefficients of *x*_1_ and *x*_2_ in the second pooling map of Interval 2). We refer to the first as the ‘disequilibrium’ and the second as the ‘conservation’ stages of relaxation for each reaction. These general characteristics can help understand the dynamic responses of more complex reaction networks.

### Toy Model 2

The second case study, we refer to as Toy Model 2, involves the same pathway as in Toy Model 1, but with energy coupling (Fig. 5). It converts a high-energy substrate (Metabolite 1) into a low-energy product (Metabolite 2). The released energy is then utilized to convert a low-energy cofactor (Metabolite 6) into its high-energy counterpart (Metabolite 5). The high-energy cofactor drives an uphill step at the beginning (Reaction 2) and is recovered in a downhill step at the end (Reaction 4). Thus, it serves a similar metabolic function to glycolysis. The cofactors are exchanged with the extracellular environment depending on their concentration gradient across the membrane, and so are the substrate and product. As in Toy Model 1, Reactions 1 and 5 are the rate-limiting steps. However, the boundary reactions for the cofactors (Reactions 6 and 7) have the largest rates. Unlike Toy Model 1, the mass-balance equations for this model are nonlinear due to the bilinearity of the mass-action rates for the cofactor-coupled reactions. In this case study, an interplay between the rate-limiting steps, cofactor exchange reactions, and cofactor-coupled intracellular reactions shape the structure of concentration pools.

**Figure 5:**
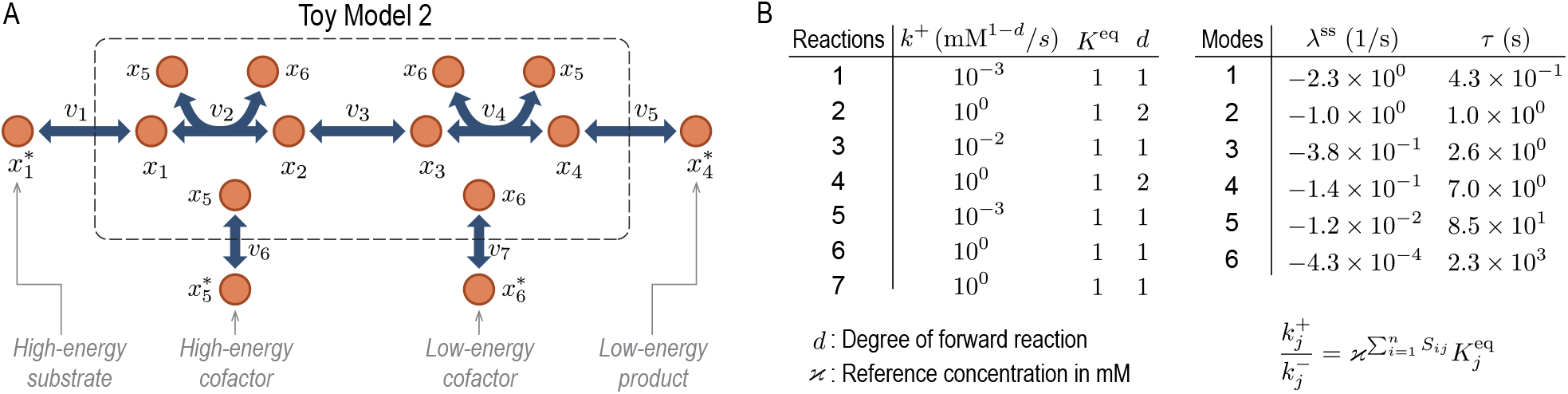
A toy model with energy coupling, where a high-energy substrate (Metabolite 1) is converted to a low-energy product (Metabolite 4). The energy released in the main pathway fuels the production a high-energy cofactor (Metabolite 5) from its low-energy counterpart (Metabolite 6). (A) Network map. (B) Rate constants and timescales. The superscript ^*∗*^ indicates metabolite concentration in the extracellular environment.

We analyzed the timescale hierarchy of Toy Model 2 by examining its dynamic response to concentration perturbations (Fig. 6A). During the relaxation of the ensuing concentration and flux disturbances, DMA identified eight time intervals associated with distinct timescales and concentration pools (Fig. 6B,C). In this model, besides reaction rates, energy coupling also influences the chronology of reactions by linking the equilibration of the cofactor-driven steps. Here, the disequilibrium stages of Reactions 2 and 4 occur on the timescale *∼* 1–10 s, and their conservation stages occur on the timescale *∼* 10–30 s. Once the concentration and flux disturbances of these reactions have relaxed, *x*_1_ and *x*_3_ form two coherent structures with *x*_2_ and *x*_4_, respectively, giving rise to two aggregate variables *x*_12_ := *x*_1_ + 2*x*_2_ and *x*_34_ := 2*x*_3_ + *x*_4_. Next, the disequilibrium stage of Reaction 3 occurs on the timescale *∼* 100–1000 s followed by a conservation stage that persists until the steady state has been reached. During this conservation stage, *x*_12_ and *x*_34_ pool together, forming a larger coherent structure. Accordingly, the slowest transients are characterized by *x*_1–4_ pooling together into a single aggregate metabolite with coefficients (1, 2, 2, 1). As the steady state is approached, the concentration of this aggregate metabolite is controlled by Reactions 1 and 5.

**Figure 6:**
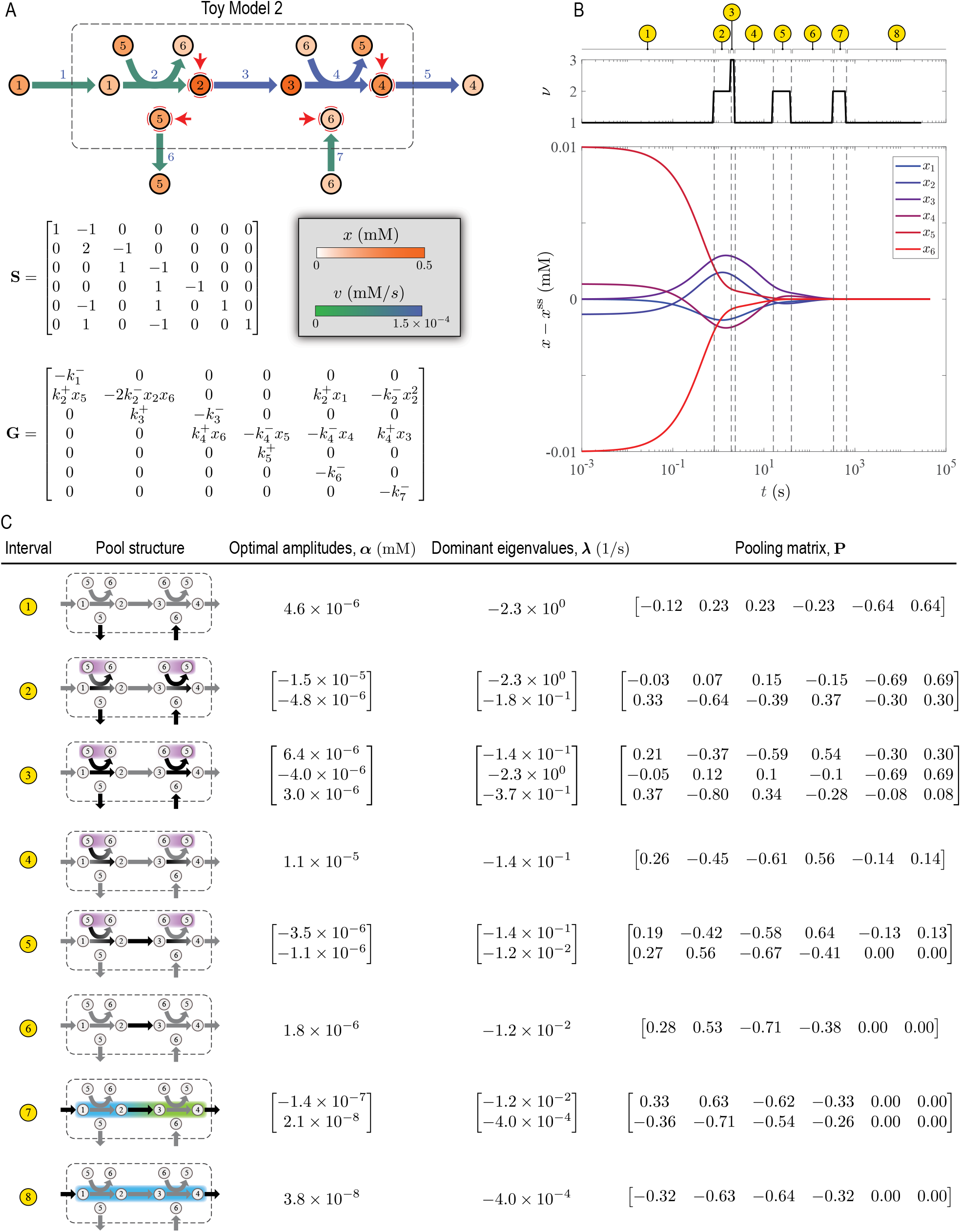
Dynamic Mode Analysis of Toy Model 2 described in Fig. 5. (A) Visualization of the steady state solution in a network map. Red arrows indicate metabolites the concentration of which is perturbed. (B) Dynamic response to concentration perturbations. Deviations from the steady state *x− x*^ss^ and dimensionality *ν* are plotted along the dynamic trajectory. Dashed lines indicate a transition between two time periods with distinct exponential decay modes, partitioning the overall relaxation time into eight characteristic time intervals. (C) Optimal amplitudes, dominant eigenvalues, and pooling matrix for each time interval identified in (B). Black arrows in pool structures indicate parts of the network that are affected the most by the initial perturbation in the corresponding time interval. Gradient and uniformly colored envelops enclose metabolites the concentration trajectory of which are negatively and positively correlated in the corresponding time interval, respectively.

Finally, we highlight the important role of cofactors in the dynamics of pathways with energy coupling. In this case study, because the cofactor exchange reactions are the fastest in the network, they control the short-term responses (*t* ≲ 30 s), only influencing the disequilibrium and conservation stages of Reactions 2 and 4. Upon concentration perturbation, *x*_5_ and *x*_6_ become negatively correlated, forming a coherent structure almost immediately (*t* ≲ 1 s). As mentioned above, the long-term responses of Toy Model 2 are mostly controlled by Reactions 1 and 5 near the steady state with minimal effect from the cofactors.

### Glycolysis

Glycolysis is a central energy-conversion pathway in biology [21]. It converts a high-energy substrate (glucose) into low-energy products (pyruvate and lactate). The energy released in this process is then used to produce high-energy cofactors (ATP and NADH). To generate a sufficient amount of high-energy phosphate bonds, it also imports a high-energy cofactor (inorganic phosphate). A key step of glycolysis is catalyzed by fructose-bisphosphate aldolase (FBP), splitting fructose 1,6-bisphosphate (fbp) into the triose phosphates dihydroxyacetone phosphate (dhap) and glyceraldehyde 3-phosphate (g3p). This step makes the energy stored in the chemical bonds of the pentose ring accessible for energy conversion in downstream reactions.

In the third case study, we analyzed the timescale hierarchy of the glycolytic pathway in human red blood cells (Fig. 7). We solved a mass-action kinetic model of glycolysis using the MASSpy package (model parameters are provided in Supplementary Data 1) [22] and computed its dynamic response to a perturbation in the ATP load (Fig. 7A). Several reactions in upper and lower glycolysis are coupled to ATP hydrolysis, so their relaxations are tightly linked together. These energy-coupling mechanisms impart an autocatalytic structure to the glycolytic pathway, a characteristic of which is oscillatory dynamics [23]. Interestingly, during the relaxation of the load perturbation, concentration trajectories exhibit oscillatory dynamics on timescales ranging from minutes to a day, coinciding with the average erythrocyte circulation time and circadian period, respectively [24]. We analyzed the concentration trajectories using DMA and identified eleven time intervals with distinct timescales and concentration pools (seven of which are highlighted in Fig. 7B,C). In the following, we highlight a few coherent structures associated with these timescales that were characterized previously. A complete list of the dominant eigenvalues and their respective pooling matrices for all time intervals is provided in Supplementary Data 2.

**Figure 7:**
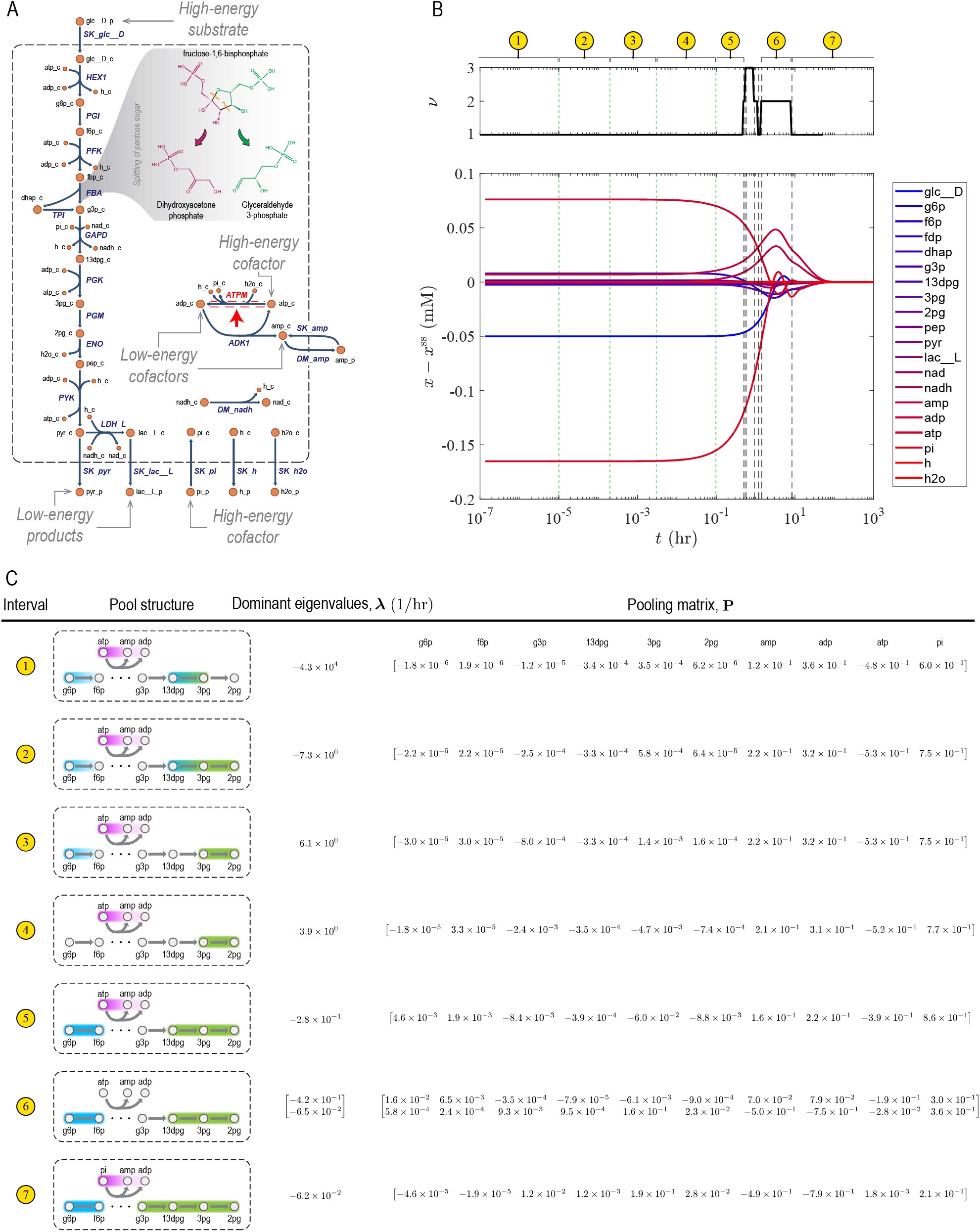
Dynamic Mode Analysis of the glycolytic pathway. (A) A network map, highlighting the energetics of substrate, products, and cofactors. The flux of the reaction indicated by the red arrow is perturbed. The retro-aldol cleavage of fructose 1,6-bisphosphate is a key step, making the energy stored in the chemical bonds of the pentose ring accessible for the production of high-energy cofactors in downstream reactions. (B) Dynamic response to flux perturbations. Deviations from the steady state *x −x*^ss^ and dimensionality *ν* are plotted along the dynamic trajectory. Dashed and dotted lines indicate a transition between two time periods with distinct exponential decay modes, partitioning the overall relaxation time into seven characteristic time intervals. Dimensionality changes across dashed but remains the same across dotted lines. (C) Dominant eigenvalues and pooling matrix for each time interval identified in (B). Gradient and uniformly colored envelops in pool structures enclose metabolites the concentration trajectory of which are negatively and positively correlated in the corresponding time interval, respectively.

The first coherent structure in this case study arises from the relaxation of phosphoglycerate mutase (PGM) on the timescale ∼0.1–100 s, where the concentrations of 3-phosphoglycerate (3pg) and 2-phosphoglycerate (2pg) become positively correlated. Next, the flux and concentration disturbances of phosphoglucoisomerase (PGI) and phosphoglycerate kinase (PGM) fully relax on the timescale ∼1–30 min. Here, the concentrations of glucose 6-phosphate (g6p) and fructose 6-phosphate (f6p) become positively correlated, forming the well-know hexose phosphate coherent structure [13]. On this timescale, the concentration of 1,3-bisphosphoglycerate (13dpg) also become correlated with those of 3pg and 2pg, leading to a larger phosphoglycerate coherent structure. On the timescale ∼10 hr, the dynamics of glyceraldehyde phosphate dehydrogenase (GAPD) relax, as a result of which glyceraldehyde 3-phosphate (g3p) merges with the phosphoglycerate coherent structure. All these coherent structures and their respective timescales are consistent with previous studies of the glycolytic dynamics using bottom-up approaches [13].

In general, coherent structures associated with slow timescales are more physiologically relevant than those forming on fast timescales [14]. Thus, we studied the two slowest pools of the glycolytic pathway forming in the last two of the eleven time intervals that DMA identified (Fig. 7B, Intervals 6 and 7), examining their respective time evolutions *π*_1_(*t*) and *π*_2_(*t*) (Fig. 8). Here, *π*_1_ and *π*_2_ are the pools associated the second slowest and slowest timescales, where 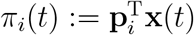 denotes the representation of pool *i* with respect to concentrations rather than concentration deviations. We found that a balance between high-energy phosphate bond and redox trafficking shapes the interconnected dynamics of upper and lower glycolysis (Fig. 8), highlighting the coupling role of the cofactors.

**Figure 8:**
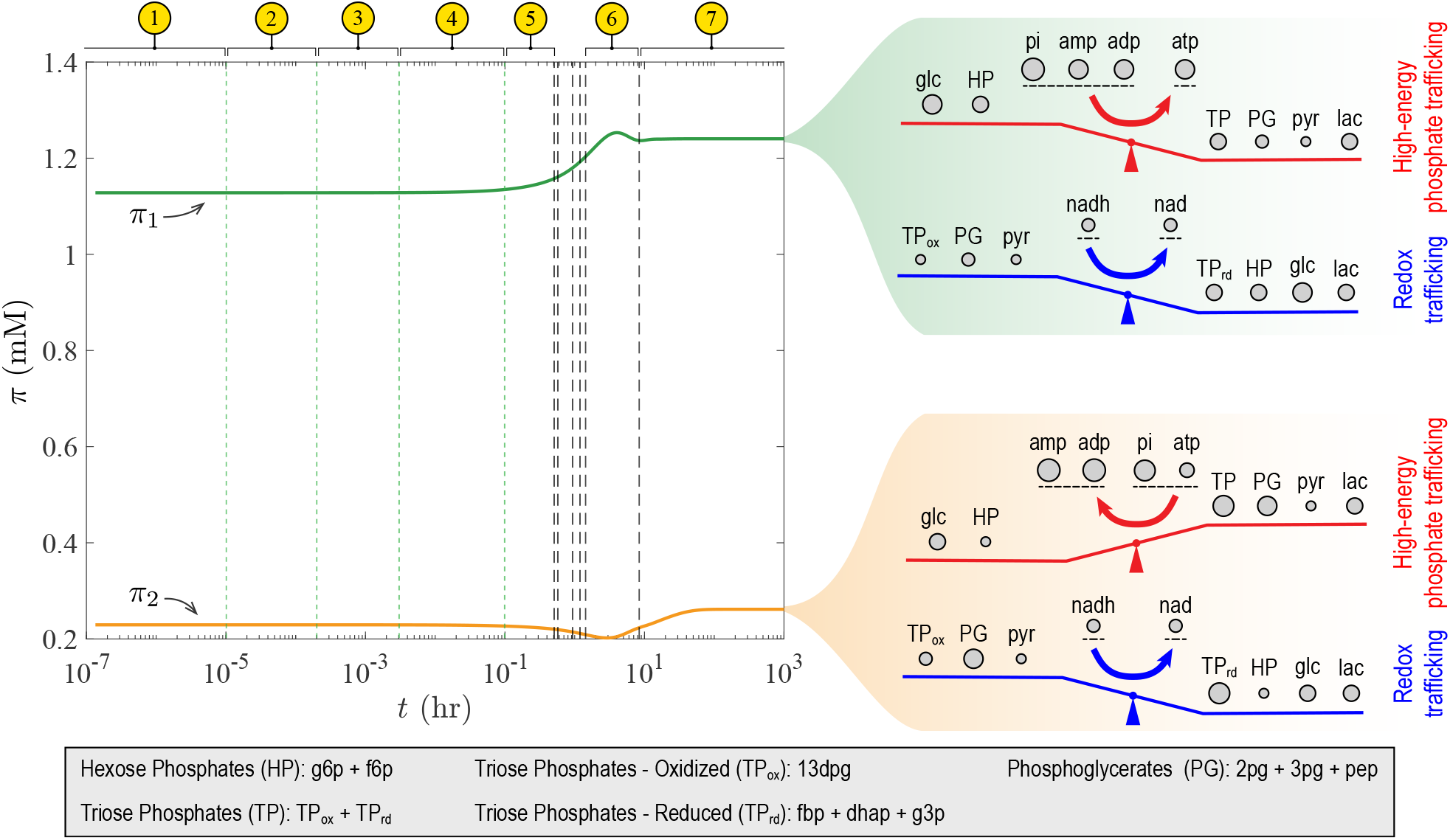
Dynamic trajectories of the slowest pools of glycolysis. The characteristic time intervals are identical to those in Fig. 7B. Note that the pools *π*_1_ and *π*_2_ are defined with respect to concentrations X rather than concentration deviations χ. Both pools are dominant in Interval 6, and *π*_2_ is the dominant pool in Interval 7. The coherent structures of *π*_1_ form near the end of Interval 6, while those of *π*_2_ form in Interval 7. The dynamics of these pools are driven by a balance between high-energy phosphate bond and redox trafficking. The circle size for each metabolite or aggregate metabolite on the right panel is proportional to its coefficient in the respective pooling map.

Finally, we note that the magnitude of the coefficients of the cofactors (AMP, ADP, ATP, and inorganic phosphate) in the pooling matrix are the largest across all timescales, implying that the glycolytic dynamics are mostly determined by the energetics of cofactor interconversions. Importantly, ATP and inorganic phosphate are the high-energy cofactors that control the dynamics on the circulation (*t* ≲ 1 min) and circadian (*t* ≳ 10 hr) timescales, respectively.

## Discussion

Living systems grow and evolve in constantly changing environments, adapting to external fluctuations through interconnected networks of chemical transformations. Understanding how these dynamics emerge from the underlying molecular processes is a major challenge of systems biology. While whole-cell kinetic models offer a thorough description of the intricate interactions within biological networks, their sheer scale often obscures the fundamental patterns that govern the system-level behavior. Using timescale decomposition techniques, we can overcome this limitation by simplifying the complex kinetics into a coarse-grained model with biologically interpretable components. This approach allows us to glean meaningful insights from the model and identify critical components that drive the dynamics at the system level.

In this paper, we introduced Dynamic Mode Analysis—a data-driven approach for timescale decomposition of chemical reaction networks. This approach characterizes concentration pools and coherent structures emerging from network dynamics using time-series data. A key component of our computational framework is an extended version of Optimal Dynamic Mode Decomposition [18, 19], which we developed to identify characteristic timescales of reaction networks. We showed that the dominant eigenvalues and eigenmodes furnished by Dynamic Mode Analysis in any time interval along dynamic trajectories are determined by two important statistical descriptors, namely the time-delayed autocorrelation and covariance. Interestingly, the former was a basis of previous top-down approaches for identification of coherent structures in reaction networks [7]. Given that timescale hierarchies often underlie the formation of dynamical patterns that lead to dimensionality reduction, our analysis provides a theoretical basis for why time-delayed autocorrelation is an effective metric for statistical analyses of coherent structures.

We validated the results of Dynamic Mode Analysis for a hypothetical pathway with analytically characterized concentration pools. We also reproduced some of the concentration pools and coherent structures of the glycolytic pathway that were determined previously using bottom-up approaches [13], further confirming the validity of our approach.

Overall, the outcomes of time-scale decomposition are more amenable to biological interpretation than the solutions of mass-balance equations. For example, our analysis of the glycolytic pathway in human red blood cells indicated two physiologically relevant characteristics: (i) Glycolysis is mostly a cofactor driven pathway, the dynamics of which are controlled by ATP and inorganic phosphate on the circulation and circadian timescales, respectively, and (ii) the slowest dynamics are dominated by a balance between high-energy phosphate bond and redox trafficking. By chronologically organizing major events along time evolutions, timescale decomposition could provide physiological insights into the dynamics of more complex biological pathways.

Quantitative models have been a chief driver of progress in biology in recent years [25–27]. Kinetics-based [1, 2, 28, 29] and constraint-based [30–35] approaches have been and will likely continue to be instrumental in these developments. Both approaches provide a mechanistic description of cellular functions, but each has its own limitations. Kinetic models are generally complex and require numerous parameters that are subject to large uncertainties, while constraint-based models are not suitable for predicting inherently dynamic phenotypes. The timescale decomposition technique presented in this study can bridge the gap between kinetic and constraint-based modeling by furnishing a systematic framework for construction of coarse-grained kinetic models [36–38] based on intrinsic timescales of biochemical reaction networks. Such reduced-order models are more tractable than whole-cell models and can capture essential characteristics of biological systems at physiologically relevant timescales.

Personalized medicine is another area where our timescale decomposition technique can play a major role. For example, when individual variations of pathological features or risk for drug side effects manifest in cellular dynamics, studying the timescale hierarchies can help identify the underlying metabolic causes [24]. Using kinetic constants as a proxy for individual genotypes in these cases, analyzing the patterns in timescales and coherent structures allows us to understand and classify disease phenotypes through the lens of timescale hierarchies.

## Appendix A1: Reciprocal Orthogonality Conditions for Toy Model 1

The physical interpretation of the reciprocal orthogonality conditions Eq. (12) is connected to the dissipative structure of chemical reaction networks. Since the transitory timescale *Ť*_1_ that we are concerned with bridges two dynamic regimes associated with a fast (*T*_1_) and slow (*T*_2_) timescale characterizing the relaxation of Reactions 2 and 3, thier respective local eigenmodes represent the equilibration direction of these reactions in the concentration space. In particular, the slow eigenmode *ϕ*_2_, which is the mode that 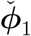 converges to when *t → T*_2_, aligns with equilibration directions of Reaction 3

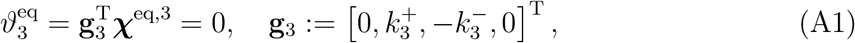

where ***χ***^eq,3^ denotes an equilibration direction. Concentration vectors that are parallel to this direction leave Reaction 3 at its steady state, while those perpendicular to this direction maximally change the flux of Reaction 3. Therefore, by requiring **p**_1_ to be orthogonal to *ϕ*_2_ in Eq. (12a), we ensure that, at any time *t* ∈ [*T*_1_, *T*_2_] along a dynamic trajectory, it points in a direction in the concentration space that maximizes the energy dissipation of Reaction 3. In contrast, the fast eigenmode *ϕ*_1_ aligns with the disequilibrium direction of Reaction 2 (see Chapter 4 of Palsson [14] for the definition of disequilibrium pools), where *χ*_1_(*t*) and *χ*_2_(*t*) are negatively correlated. Thus, by requiring 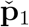 to be orthogonal to *ϕ*_1_ in Eq. (12b), we ensure that it points in a direction corresponding to the conservation of metabolites involved in Reaction 2 (see Chapter 4 of Palsson [14] for the definition of conservation pools), where *χ*_1_(*t*) and *χ*_2_(*t*) are positively correlated.

The reciprocal orthogonality conditions in Eq. (12) also have an important geometric interpretation. For a given time interval along a dynamic trajectory, dominant eigenmodes represent the principal directions in a part of the concentration space where the dynamics occur. Therefore, if the underlying reaction system in the time interval of interest is low-dimensional, the eigenmodes can be viewed as a natural basis spanning a reduced concentration space where dynamic trajectories lie. Regarding this reduced concentration space as primal space (see Fig. 1F), Eq. (12) ensures that the vector representation of pools are a natural basis of a corresponding dual space. From this standpoint, the pools *p*_1_ and 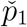 in Toy Model 1 are scalar quantities defined as the inner product of a vector in the primal with another vector in the dual space, the algebraic form of which remain invariant with respect to any linear coordinate transformation [39]. This geometric interpretation is compatible with the physical interpretation of flux-concentration duality in dissipative systems [16]. As mentioned above, the vector representation of pools are closely related to flux-disturbance relaxation or maximal energy dissipation of reactions, so they form a basis for a dual space in which to represent flux dynamics naturally. Similarly, eigenmodes constitute a basis for a primal space in which to represent concentration dynamics naturally.

## Appendix A2: Derivation of the Third and Fourth Approximate Eigenmodes of Toy Model 1

Here, we provide more details on the derivation of approximate expressions for the third and fourth eigenmodes of Toy Model 1 discussed in the main text. We note that the third eigenmode *ϕ*_3_ arises from the second transitory eigenmode 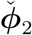 in the interval [ *Ť*_2_, *T*_3_], we may rega rd *ϕ*_3_ as the limit of 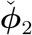 when *t → T*_3_. The second transitory regime occurs in the interval [ Ť_2_, *T*_3_] when Reaction 3 equilibrates. In this transition period, the flux deviations of Reaction 3 and 4 are of the same order, so that *ϑ*_2_ *≃ θ*^a^*ϑ*_3_ and *ϑ*_3_ *∼ ϑ*_4_ *≫ ϑ*_1_, *ϑ*_5_. As with the first transitory period, we consider a coefficient *β* such that *ϑ*_3_ = *βϑ*_4_ to quantify the transition between *T*_2_ and *T*_3_ with *β ∼ 𝒪*(1). Accordingly, the mass-balance equations simplify to

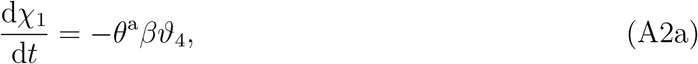

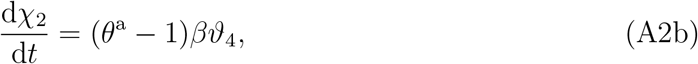

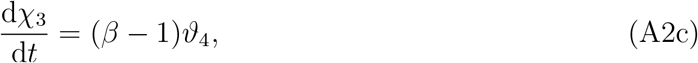

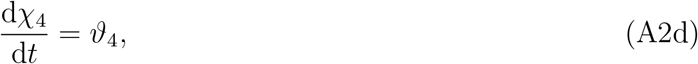

Eliminating *χ*_3_ and *χ*_4_, Eq. (A2) is expressed with respect to *ϑ*_4_ as

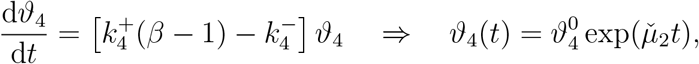

where 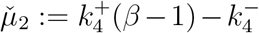, and 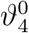 is the initial condition for the flux deviation of Reaction 4. Substituting the solution of *ϑ*_4_ back in Eq. (A2), we arrive at

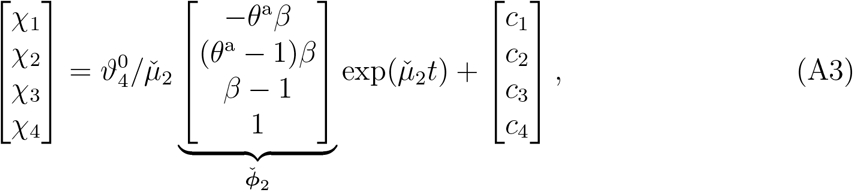

where *c*_*i*_ are the integration constants. Note that the coefficient *β* varies from a large value to its asymptotic value *β*^a^ as *t →T*_3_. This asymptotic coefficient is in turn determined from the equilibrium of Reaction 3

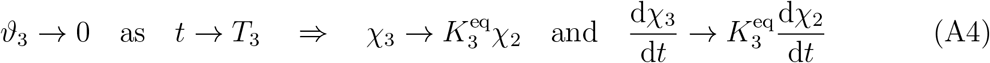

with 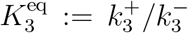 the equilibrium constant of Reaction 3. It follows from Eqs. (A4), (A2b), and (A2c) that

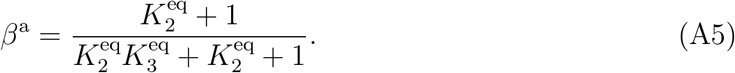

We now can express *ϕ*_3_ and *µ*_3_ as the limiting cases of 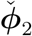 and 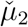

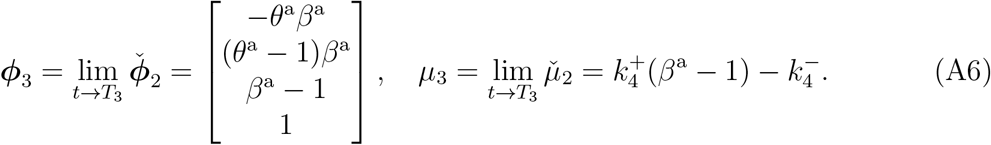

The third transitory regime occurs in the interval [*Ť*_3_, *T*_*∞*_], where *T*_*∞*_ is a sufficiently large time—referred to as the total relaxation time—by which all flux and concentration disturbances have relaxed to within a tolerable threshold. This regime is characterized by *ϑ*_2_ *≃ θ*^a^*ϑ*_3_, *ϑ*_3_ *≃ β*^a^*ϑ*_4_, and *ϑ*_4_ *∼ ϑ*_1_, *ϑ*_5_. We express the order-of-magnitude balance between the flux deviations of Reaction 4 and 5 by a coefficient *γ* such that *ϑ*_4_ = *γϑ*_5_ to describe the transitory regime between *T*_3_ and *T*_*∞*_ with *γ ∼ 𝒪*(1), simplifying the mass-balance equations to

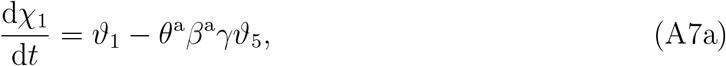

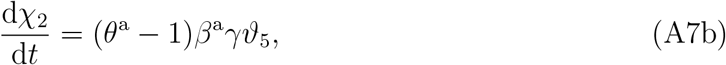

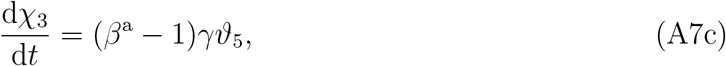

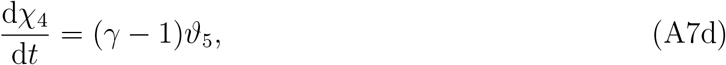

From Eq. (A7d) it follows that

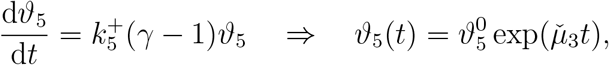

where 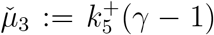, and 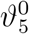 is the initial condition for the flux deviation of Reaction 5. Substituting *ϑ*_5_(*t*) from this equation in Eq. (A7a), we obtain the following solution for *ϑ*_1_(*t*)

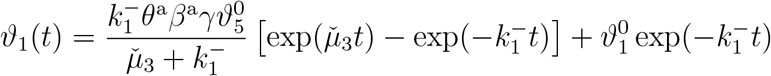

with 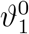 the initial condition for the flux deviation of Reaction 5. From this solution, it follows that the dynamics of *ϑ*_1_ in the third transitory regime is described by two exponential decay modes of the same order-of-magnitude (Note that 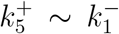 in Toy Model 1; see Fig. 2C). Our goal in the analysis of this transitory regime is to ascertain the slowest eigenmode of Toy Model 1 and describe the dynamics near its steady state. Therefore, we assume 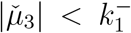 in the remainder of this section, so that the slowest timescale is characterized by 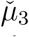. A similar analysis can be performed for the opposite case. Substituting the solutions of *ϑ*_1_ and *ϑ*_5_ in Eq. (A7) and neglecting the slower mode associated with exp 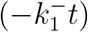, we obtain the dynamic trajectory of concentration deviations

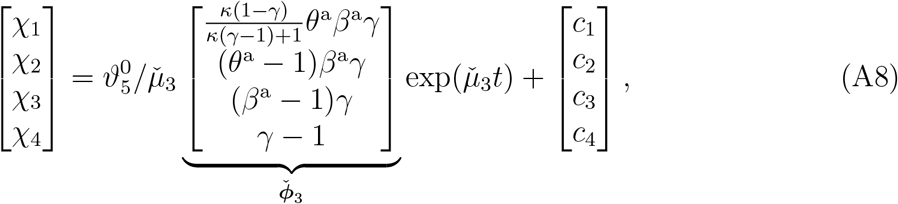

where 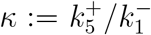. Since the flux deviations of Reaction 1 becomes non-negligible in this regime, the equilibration of Reaction 2 and 3 is disturbed once again on the third transitory timescale *Ť*_3_. Therefore, the asymptotic coefficients *θ*^a^ and *β*^a^ change during the secondary relaxation of Reaction 2 and 3 as *t →T*_*∞*_. We determine the asymptotic value of *θ* in the third transitory regime from Eqs. (6), (A7a), and (A7b)

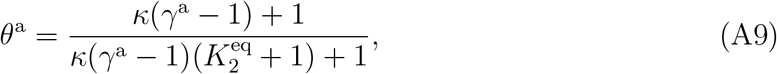

and of *β* from Eqs. (A4), (A7b), and (A7c)

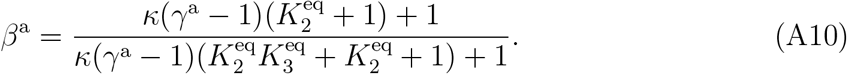

The coefficient *γ* varies from a large value to its asymptotic value *γ*^a^ as *t →T*_*∞*_, which is ascertained from the equilibrium of Reaction 4

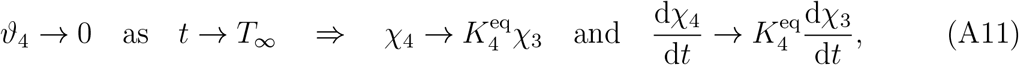

where 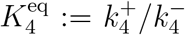 is the equilibrium constant of Reaction 4. From Eqs. (A11), (A7c), and (A7d), we find that

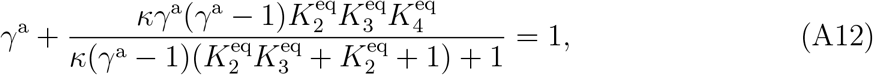

which is a quadratic polynomial to be solved with respect to *γ*^a^. As with previous cases, *ϕ*_4_ and *µ*_4_ are the limiting cases of 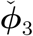 and 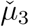, so that

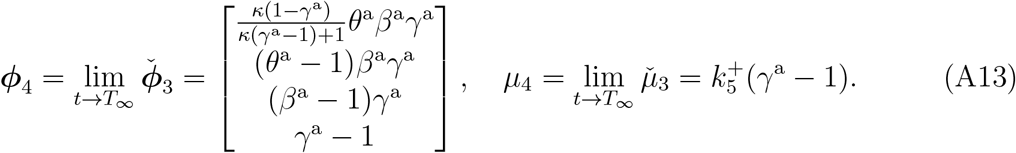

## Appendix A3: Inhomogeneity of evolution equations for local concentration devaiations and temporal grid size

We show that the discrete form of evolution equations for local concentration deviations Eq. (34) becomes homogeneous asymptotically for vanishingly small temporal grid size, that is ***ω*** *→* **0** as Δ*t →* 0. We start from the definition of ***ω*** in Eq. (34)

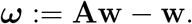

Substituting the definitions of **A** and **w** in the main text (see Section “Dynamic Mode Analysis”) in this equation, we have

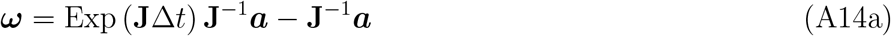

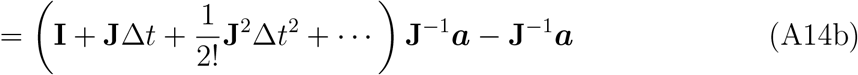

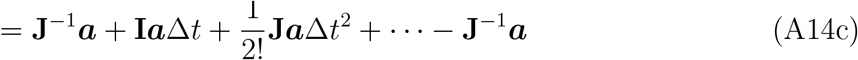

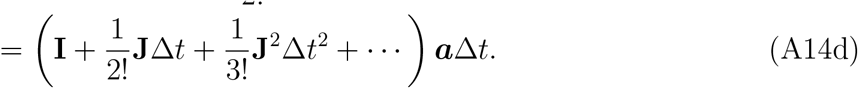

In Eq. (A14d), the expression enclosed in parentheses is a matrix with a nonzero norm. Moreover, the constant vector a is generally nonzero in all sliding time windows, and it only approaches zero asymptotically as dynamic trajectories near a steady-state solution. It follows that, the inhomogeneous term ***ω*** approaches zero asymptotically in time windows away from the steady state only if an arbitrarily small temporal grid size Δ*t* is chosen.

## Appendix A4: Relationship between time-delayed autocorrelation, covariance, and the local Jacobian spectra

Here, we prove that 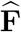 and 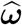 in Eq. (38) are the solution of the optimization problem Eq. (37). We begin by expanding the objective

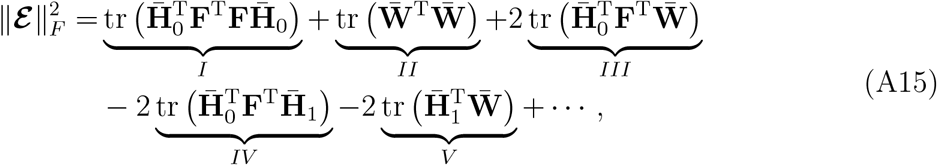

where 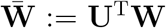, and the dots represent constant terms that do not affect the optimal solution of Eq. (37). As stated in the main text, **U** can be regarded as a transformation, mapping matrices and vectors from the space of proper orthogonal modes into the original concentration space, so we may write 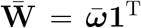 with 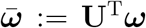. In the following, we further expand each term in Eq. (A15) to derive expressions that can be readily recast into a standard-form quadratic program.

Starting from the first term

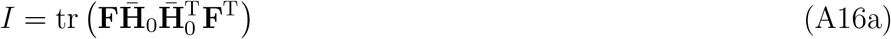

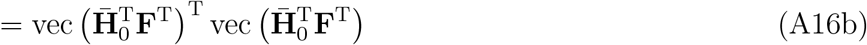

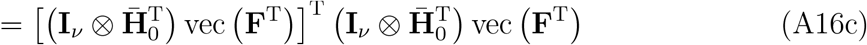

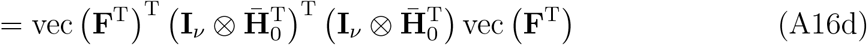

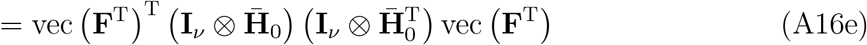

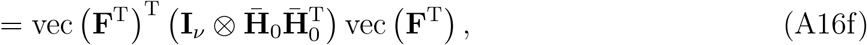

where **I**_*ν*_ denotes a *ν ×ν* identity matrix, and vec (·) vectorizes a matrix by stacking its columns on top of one another. Here, we used the cyclic property of the trace in Eq. (A16a) and applied the identity vec 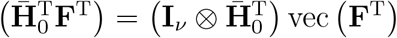 in Eq. (A16c). Expanding the second term, we arrive at

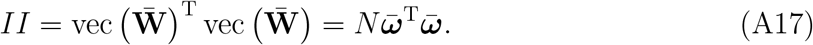

The third term is written

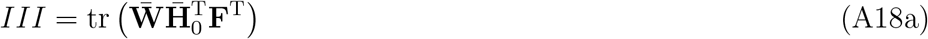

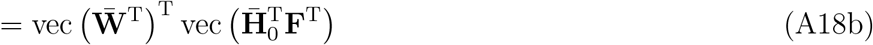

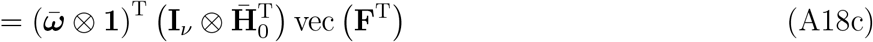

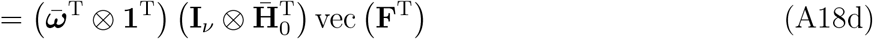

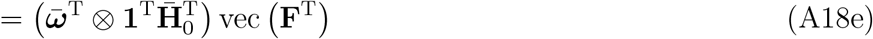

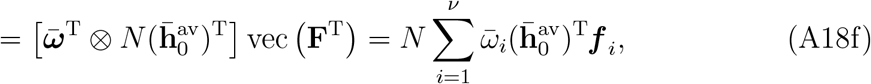

where ***f*** _*i*_ is the *i*th column of **F**^T^. Again, we used the cyclic property of the trace in Eq. (A18a) and the identity vec 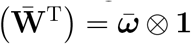 in Eq. (A18c). Expanding the fourth term, we have

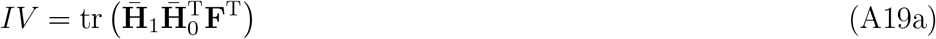

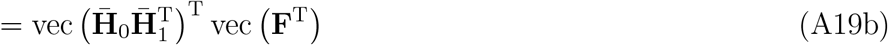

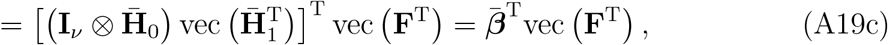

where

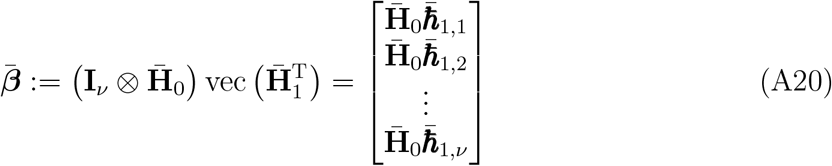

with 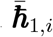 denoting the *i*th column of 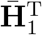. The fifth term is expanded similarly

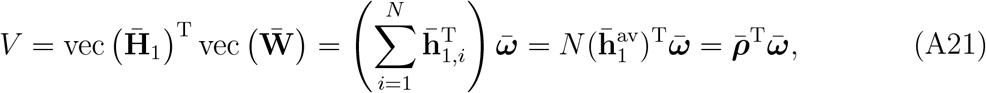

where 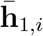 denotes the *i*th column of 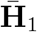, and 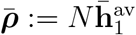.

We now can rewrite the optimization problem Eq. (37) as a quadratic program in standard form by substituting the five terms *I*–*V* from Eqs. (A16)–(A21) in Eq. (A15)

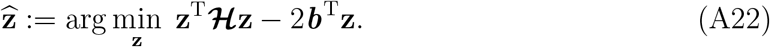

Here,

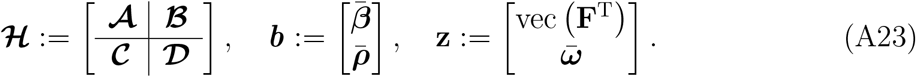

with

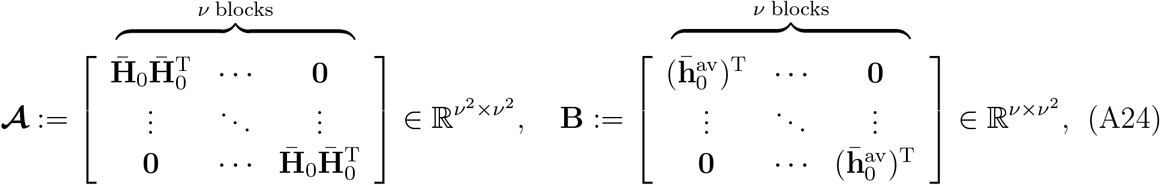

***ℬ***:= *N* **B**^T^, ***𝒞***:= *N* **B**, and ***𝒟***:= *N* **I**_*ν*_. The solution of the minimization problem Eq. (A22) is ascertained from the Karush-Kuhn-Tucker conditions [40]

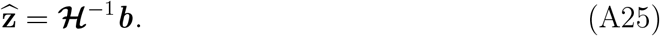

To derive compact-from expressions for 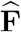 and 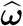, the Hessian ***ℋ*** is to be inverted explicitly with respect to its elements. Since the Hessian in Eq. (A23) is a block matrix, we derive its block inverse

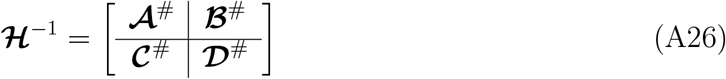

using the Schur complement. With respect to the matrix blocks in Eq. (A23), the Schur complement of ***ℋ***is written

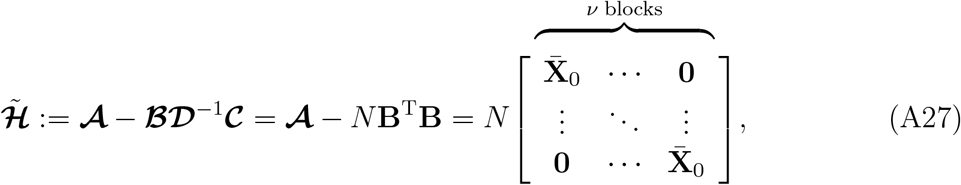

where 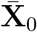 is the covariance of time-series data 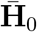 defined in Eq. (39b). We can now express the individual blocks in Eq. (A26) in terms of the Schur complement 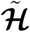

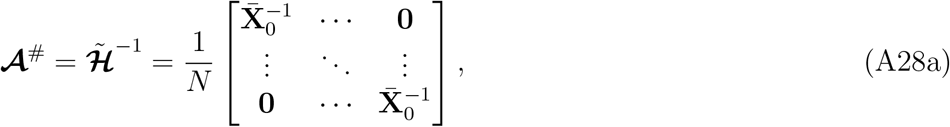

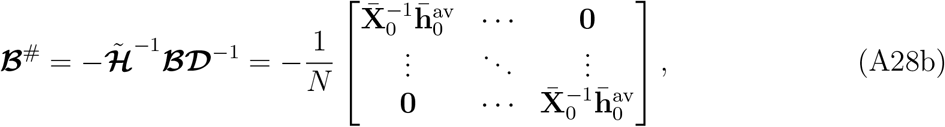

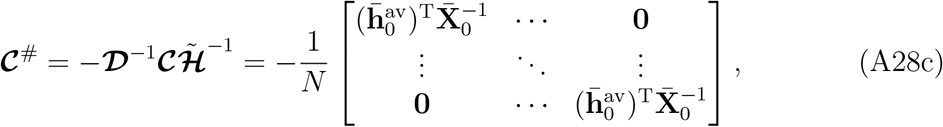

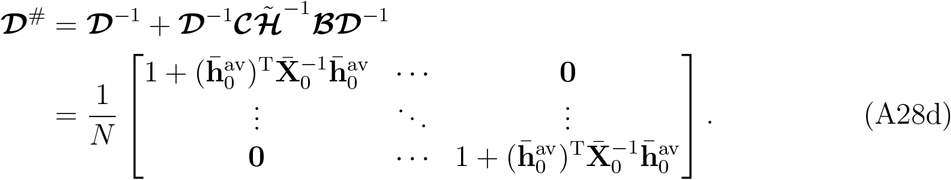

Accordingly, the two parts of the optimal solution 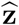 are ascertained separated by plugging ***ℋ***^*−*1^ from Eq. (A26) into Eq. (A25)

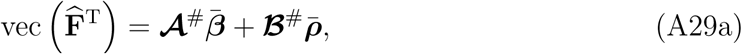

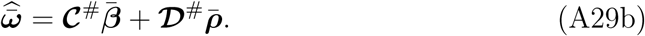

Substituting the inverse blocks from Eqs. (A28a) and (A28b) in Eq. (A29a), we obtain the first part

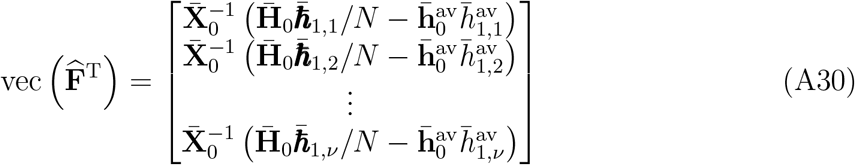

with 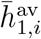 denoting the *i*th component of 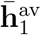. From Eq. (A30), one can verify that the optimal solution in matrix form is

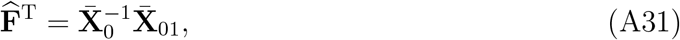

where

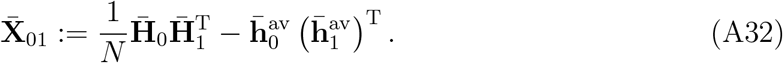

Since 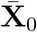 is symmetric and 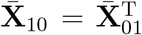, the optimal solution 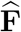 given in Eq. (38a) in the main text follows from Eq. (A31). Similarly, the second part of the optimal solution is obtained by substituting the inverse blocks from Eqs. (A28c) and (A28d) in Eq. (A29b)

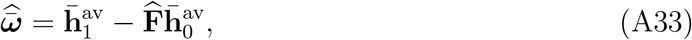

from which the optimal solution 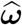 given in Eq. (38b)is derived using the relationship 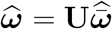.

## Author Contributions

Conceptualization, A.A. and B.O.P.; Methodology, A.A.; Validation A.A.; Formal Analysis A.A.; Investigation, A.A. and Z.B.H.; Writing – Original Draft, A.A.; Writing – Review & Editing, A.A., Z.B.H., and B.O.P.; Funding Acquisition, B.O.P.; Resources, B.O.P.; Supervision, B.O.P.

## Data Availability

All data generated or analyzed during this study are included in this published article and its Supplementary Information files.

## Code Availability

All the codes, their description, and data used for analysis are provided in Supplementary Information files.

## Acknowledgments

This work was funded by the Novo Nordisk Foundation (Grant Number NNF10CC1016517) and the National Institutes of Health (Grant Number GM057089).

## References

1. Karr, J.R., Sanghvi, J.C., Macklin, D.N., Gutschow, M.V., Jacobs, J.M., Bolival, B., Assad-Garcia, N., Glass, J.I., Covert, M.W.. A whole-cell computational model predicts phenotype from genotype. Cell 2012;150(2):389–401. doi: 10.1016/j.cell.2012.05.044.

2. Macklin, D.N., Ahn-Horst, T.A., Choi, H., Ruggero, N.A., Carrera, J., Mason, J.C., Sun, G., Agmon, E., DeFelice, M.M., Maayan, I., et al. Simultaneous cross-evaluation of heterogeneous E. coli datasets via mechanistic simulation. Science 2020;369(6502). doi:10.1126/science.aav3751.

3. Palsson, B.O., Joshi, A.. On the dynamic order of structured escherichia coli growth models. Biotechnol Bioeng 1987;29(6):789–792. doi:10.1002/bit.260290623.

4. Heinrich, R., Sonntag, I.. Dynamics of non-linear biochemical systems and the evolutionary significance of time hierarchy. Biosystems 1982;15(4):301–316. doi: 10.1016/0303-2647(82)90045-4.

5. Stelling, J., Sauer, U., Szallasi, Z., Doyle III, F.J., Doyle, J.. Robustness of cellular functions. Cell 2004;118(6):675–685. doi:10.1016/j.cell.2004.09.008.

6. Hintze, A., Adami, C.. Evolution of complex modular biological networks. PLoS Comput Biol 2008;4(2):e23. doi:10.1371/journal.pcbi.0040023.

7. Arkin, A., Ross, J.. Statistical construction of chemical reaction mechanisms from measured time-series. J Phys Chem 1995;99(3):970–979. doi:10.1021/j100003a020.

8. Arkin, A., Shen, P., Ross, J.. A test case of correlation metric construction of a reaction pathway from measurements. Science 1997;277(5330):1275–1279. doi: 10.1126/science.277.5330.1275.

9. Kauffman, K.J., Pajerowski, J.D., Jamshidi, N., Palsson, B.O., Edwards, J.S.. Description and analysis of metabolic connectivity and dynamics in the human red blood cell. Biophys J 2002;83(2):646–662. doi:10.1016/s0006-3495(02)75198-9.

10. Jamshidi, N., Palsson, B.O.. Top-down analysis of temporal hierarchy in biochemical reaction networks. PLoS Comput Biol 2008;4(9):e1000177. doi: 10.1371/journal.pcbi.1000177.

11. Schmid, P.J.. Dynamic mode decomposition of numerical and experimental data. J Fluid Mech 2010;656:5–28. doi:10.1017/s0022112010001217.

12. Ben-Israel, A., Greville, T.N.E.. Generalized inverses: Theory and applications; vol. 15. Springer Science & Business Media; 2003.

13. Jamshidi, N., Palsson, B.O.. Formulating genome-scale kinetic models in the postgenome era. Mol Syst Biol 2008;4(1):171. doi:10.1038/msb.2008.8.

14. Palsson, B.O.. Systems biology: Simulation of dynamic network states. Cambridge University Press; 2011.

15. Hyvärinen, A., Oja, E.. Independent component analysis: Algorithms and applications. Neural Netw 2000;13(4-5):411–430. doi:10.1016/s0893-6080(00)00026-5.

16. Warren, P.B., Jones, J.L.. Duality, thermodynamics, and the linear programming problem in constraint-based models of metabolism. Phys Rev Lett 2007;99(10):108101. doi:10.1103/physrevlett.99.108101.

17. Ivancevic, V.G., Ivancevic, T.T.. Applied differential geometry: A modern introduction. World Scientific; 2007.

18. Wynn, A., Pearson, D.S., Ganapathisubramani, B., Goulart, P.J.. Optimal mode decomposition for unsteady flows. J Fluid Mech 2013;733:473–503. doi: 10.1017/jfm.2013.426.

19. Jovanović, M.R., Schmid, P.J., Nichols, J.W.. Sparsity-promoting dynamic mode decomposition. Phys Fluids 2014;26(2). doi:10.1063/1.4863670.

20. Edwards, W.S., Tuckerman, L.S., Friesner, R.A., Sorensen, D.C.. Krylov methods for the incompressible navier-stokes equations. J Comput Phys 1994;110(1):82–102. doi:10.1006/jcph.1994.1007.

21. Berg, J.M., Tymoczko, J.L., Stryer, L.. Biochemistry. Fifth ed.; Freeman: New York; 2002.

22. Haiman, Z.B., Zielinski, D.C., Koike, Y., Yurkovich, J.T., Palsson, B.O. Masspy: Building, simulating, and visualizing dynamic biological models in python using mass action kinetics. PLoS Comput Biol 2021;17(1):e1008208. doi: 10.1371/journal.pcbi.1008208.

23. Chandra, F.A., Buzi, G., Doyle, J.C.. Glycolytic oscillations and limits on robust efficiency. Science 2011;333(6039):187–192. doi:10.1126/science.1200705.

24. Bordbar, A., McCloskey, D., Zielinski, D.C., Sonnenschein, N., Jamshidi, N., Palsson, B.O.. Personalized whole-cell kinetic models of metabolism for discovery in genomics and pharmacodynamics. Cell Syst 2015;1(4):283–292. doi: 10.1016/j.cels.2015.10.003.

25. Mogilner, A., Wollman, R., Marshall, W.F.. Quantitative modeling in cell biology: What is it good for? Dev Cell 2006;11(3):279–287. doi:10.1016/j.devcel.2006.08.004.

26. McCloskey, D., Palsson, B.O., Feist, A.M.. Basic and applied uses of genome-scale metabolic network reconstructions of escherichia coli. Mol Syst Biol 2013;9(1):661. doi:10.1038/msb.2013.18.

27. Del Sol, A., Jung, S.. The importance of computational modeling in stem cell research. Trends Biotechnol 2021;39(2):126–136. doi:10.1016/j.tibtech.2020.07.006.

28. Joshi, A., Palsson, B.O.. Metabolic dynamics in the human red cell: Part i—a comprehensive kinetic model. J Theor Biol 1989;141(4):515–528. doi:10.1016/S0022-5193(89)80233-4.

29. Berndt, N., Bulik, S., Wallach, I., Wünsch, T., König, M., Stockmann, M., Meierhofer, D., Holzhütter, H.G.. Hepatokin1 is a biochemistry-based model of liver metabolism for applications in medicine and pharmacology. Nat Commun 2018;9(1):2386. doi:10.1038/s41467-018-04720-9.

30. Segre, D., Vitkup, D., Church, G.M.. Analysis of optimality in natural and perturbed metabolic networks. Proc Natl Acad Sci 2002;99(23):15112–15117. doi: 10.1073/pnas.232349399.

31. Beg, Q.K., Vazquez, A., Ernst, J., de Menezes, M.A., Bar-Joseph, Z., Barabási, A.L., Oltvai, Z.N.. Intracellular crowding defines the mode and sequence of substrate uptake by escherichia coli and constrains its metabolic activity. Proc Natl Acad Sci 2007;104(31):12663–12668. doi:10.1073/pnas.0609845104.

32. Henry, C.S., Broadbelt, L.J., Hatzimanikatis, V.. Thermodynamics-based metabolic flux analysis. Biophys J 2007;92(5):1792–1805. doi:10.1529/biophysj.106.093138.

33. Lerman, J.A., Hyduke, D.R., Latif, H., Portnoy, V.A., Lewis, N.E., Orth, J.D., Schrimpe-Rutledge, A.C., Smith, R.D., Adkins, J.N., Zengler, K., Palsson, B.O.. In silico method for modelling metabolism and gene product expression at genome scale. Nat Commun 2012;3(1):1–10. doi:10.1038/ncomms1928.

34. Akbari, A., Palsson, B.O.. Scalable computation of intracellular metabolite concentrations. Comput Chem Eng 2021;145:107164. doi:10.1016/j.compchemeng.2020.107164.

35. Akbari, A., Yurkovich, J.T., Zielinski, D.C., Palsson, B.O.. The quantitative metabolome is shaped by abiotic constraints. Nat Commun 2021;12(1):1–19. doi: 10.1038/s41467-021-23214-9.

36. Basan, M., Hui, S., Okano, H., Zhang, Z., Shen, Y., Williamson, J.R., Hwa, T.. Overflow metabolism in escherichia coli results from efficient proteome allocation. Nature 2015;528(7580):99–104. doi:10.1038/nature15765.

37. Erickson, D.W., Schink, S.J., Patsalo, V., Williamson, J.R., Gerland, U., Hwa, T.. A global resource allocation strategy governs growth transition kinetics of escherichia coli. Nature 2017;551(7678):119–123. doi:10.1038/nature24299.

38. Avanzini, F., Freitas, N., Esposito, M.. Circuit theory for chemical reaction networks. Phys Rev X 2023;13(2):021041. doi:10.1103/physrevx.13.021041.

39. Wrede, R.C.. Introduction to vector and tensor analysis. Courier Corporation; 2013.

40. Nocedal, J., Wright, S.. Numerical optimization. Springer Science & Business Media; 2006.

